# Massively parallel characterization of transcriptional regulatory elements in three diverse human cell types

**DOI:** 10.1101/2023.03.05.531189

**Authors:** Vikram Agarwal, Fumitaka Inoue, Max Schubach, Beth K. Martin, Pyaree Mohan Dash, Zicong Zhang, Ajuni Sohota, William Stafford Noble, Galip Gürkan Yardimci, Martin Kircher, Jay Shendure, Nadav Ahituv

**Affiliations:** Department of Genome Sciences, University of Washington, Seattle, WA 98195, USA; mRNA Center of Excellence, Sanofi Pasteur Inc., Waltham, MA 02451, USA; Department of Bioengineering and Therapeutic Sciences, University of California San Francisco, San Francisco, CA 94158, USA; Institute for Human Genetics, University of California San Francisco, San Francisco, CA 94158, USA; Institute for the Advanced Study of Human Biology (WPI-ASHBi), Kyoto University, Kyoto, Japan; Berlin Institute of Health of Health at Charité - Universitätsmedizin Berlin, 10178, Berlin, Germany; Paul G. Allen School of Computer Science and Engineering, University of Washington, Seattle, WA, USA; Knight Cancer Institute, Oregon Health and Science University, Portland, OR, USA; Cancer Early Detection Advanced Research Center, Oregon Health and Science University, Portland, OR, USA; Institute of Human Genetics, University Medical Center Schleswig-Holstein, University of Lübeck, Lübeck, Germany; Howard Hughes Medical Institute, Seattle, WA 98195, USA; Brotman Baty Institute for Precision Medicine, University of Washington, Seattle, WA 98195, USA; Allen Center for Cell Lineage Tracing, University of Washington, Seattle, WA 98195, USA

**Author notes:** Correspondence to Jay Shendure and Nadav Ahituv. These authors contributed equally to this work (Vikram Agarwal, Fumitaka Inoue).

## Abstract

The human genome contains millions of candidate *cis*-regulatory elements (CREs) with cell-type-specific activities that shape both health and myriad disease states. However, we lack a functional understanding of the sequence features that control the activity and cell-type-specific features of these CREs. Here, we used lentivirus-based massively parallel reporter assays (lentiMPRAs) to test the regulatory activity of over 680,000 sequences, representing a nearly comprehensive set of all annotated CREs among three cell types (HepG2, K562, and WTC11), finding 41.7% to be functional. By testing sequences in both orientations, we find promoters to have significant strand orientation effects. We also observe that their 200 nucleotide cores function as non-cell-type-specific ‘on switches’ providing similar expression levels to their associated gene. In contrast, enhancers have weaker orientation effects, but increased tissue-specific characteristics. Utilizing our lentiMPRA data, we develop sequence-based models to predict CRE function with high accuracy and delineate regulatory motifs. Testing an additional lentiMPRA library encompassing 60,000 CREs in all three cell types, we further identified factors that determine cell-type specificity. Collectively, our work provides an exhaustive catalog of functional CREs in three widely used cell lines, and showcases how large-scale functional measurements can be used to dissect regulatory grammar.

## INTRODUCTION

Sequence variation in gene regulatory elements is a major cause of human disease^1–3^. For example, the overwhelming majority of genome-wide association studies (GWAS) for common disease unambiguously implicate noncoding haplotypes that contain distal gene regulatory elements, such as enhancers, in numerous disorders^4–7^. However, identifying, characterizing, and/or predicting the functional effects of nucleotide variation in gene regulatory elements remains challenging for the following reasons: 1) the lack of a ‘regulatory code’ through which the molecular consequences of a given noncoding variant might be accurately predicted; 2) the lack of a comprehensive delineation of the likely millions of regulatory elements in the human genome; 3) the difficulty of assigning regulatory elements to the genes that they regulate; 4) the cell-type specificity and/or association of regulatory elements, as well as noncoding variant effects, with developmentally or homeostatically transient cell states; 5) the minute effect sizes of many phenotypically relevant variants; and 6) the potential redundancy and/or cooperativity between regulatory elements.

The emergence of genome-scale biochemical technologies to globally catalog regions of open chromatin (DNase-seq, ATAC-seq), transcription factor (TF) binding, histone modifications (ChIP-seq or CUT&RUN) and mRNA expression levels (RNA-seq) has provided a framework to investigate gene regulatory and transcriptional landscapes in hundreds of cell types^8, 9^. These efforts have led to the discovery of millions of CREs in the human genome. However, this type of approach has some inherent limitations: 1) These types of data are descriptive, providing the potential for a sequence to be a CRE, but not a functional assay or readout to actually test this; 2) These descriptive data do not tell us which gene(s) are regulated by any given element, the magnitude of activity (*i.e.*, the level of transcriptional activation conferred by a promoter or enhancer), nor the impact of genetic variation on that activity (*e.g.*, the effect of a single nucleotide variant within a predicted enhancer that could underlie a proximally located GWAS hit or disease association); 3) The resolution of descriptive methods tends to be coarse. For example, it is unclear how the boundaries of ChIP-seq-defined binding peaks relate to the true boundaries of the corresponding CRE with respect to its functional activity. Furthermore, such assays provide little insight into how individual regulatory elements are internally organized to mediate a specific pattern of expression, and are intrinsically limited by the availability of specific antibodies. Other assays such as DNase-seq may provide greater resolution, but are nevertheless fundamentally descriptive. For example, DNase I hypersensitivity will not distinguish between spurious and functional binding events of a given TF, nor does it inform the functional identity of a regulatory element (*e.g.*, enhancer vs. promoter), its magnitude of activity, or the gene(s) that it regulates.

Massively parallel reporter assays (MPRAs) overcome many of these limitations by testing thousands of sequences/variants for their regulatory activity in a multiplex fashion^10^. By linking thousands of sequences to a transcribed sequence-based barcode and measuring their transcriptional activity via RNA-seq, normalized for its cellular integration by sequencing the corresponding DNA barcode (via DNA-seq), MPRAs provide a functional and quantitative measurement for thousands of CRE sequences/variants independent of their location and epigenetic context. Previous work has utilized self-transcribing active regulatory region sequencing (STARR-seq) to test a large number of sequences for regulatory activity in human cells^11–14^. This includes, for example, the testing of various TF binding sequences (TFBSs) in 49 bp chunks, 500 bp human genome random fragments, 170 bp synthetic random sequences, and 150 bp synthetic enhancer–promoter combinations in human colon carcinoma cells (GP5d)^14^. Another example involved the deployment of whole-genome STARR-seq in human prostate cancer cells (LNCaP)^12^. However, these assays utilize transfection to introduce a large number of sequences, providing an episomal (‘out of genome’) readout, and only work in a limited number of established cell types that can be robustly transfected and grown in large quantities. To address these and other limitations of episomal MPRAs, we previously developed a lentivirus-based MPRA (lentiMPRA) that enables reproducibility and multiplexability, the ability to carry out these assays in ‘hard to transfect cell lines’, and provides an ‘in genome’ readout whose results are more strongly correlated with ENCODE annotations and sequence-based models^15^. However, this method had been limited in terms of the number of sequences/variants that could be tested in a single experiment (*i.e.*, up to 14,000)^16, 17^.

Here, we further develop lentiMPRA and confirm the reproducibility and reliability of this technology to test over 200,000 sequences in a single experiment, which covers the majority of CREs of any given human cell type^18^. We utilize our substantially expanded MPRA data in three major ENCODE cell types, human hepatocytes (HepG2), lymphoblasts (K562), and induced pluripotent stem cells (iPSCs; WTC11), to examine the relative orientation dependence of promoters and enhancers. Furthermore, we test 60,000 sequences in all three cell lines. Utilizing our data, we characterize the activity effect of a core promoter region and train models that can characterize regulatory activity in both promoters and enhancers. We identify both biochemical and sequence-based features that are associated with cell-type-specific activity and provide a catalog of thousands of functional CREs, a unique and nearly comprehensive dataset that advances our understanding of genotype-to-phenotype associations in gene regulatory sequences.

## RESULTS

### Optimization of lentiMPRA technology

To scale up lentiMPRA^15, 19^, we revised our established protocol to add random barcodes to the assayed sequences during the library amplification step^16, 17, 20^. This framework consisted of: i) ordering an Agilent library of oligonucleotides, ii) adding a minimal promoter and random barcodes through two sequential PCR steps, and iii) cloning these elements into the pLS-SceI vector (**Extended Data Fig. 1a**). Subsequently, element-barcode associations were reconstructed through sequencing (**Extended Data Fig. 1b**) and analyzed with MPRAflow^20^. To evaluate the robustness of this revised lentiMPRA approach, we designed two pilot libraries (**Extended Data Fig. 2a**). The first library encompassed 9,372 elements in HepG2 cells, and consisted of: i) 9,172 putative enhancers, centered at DNase hypersensitivity peaks which did not overlap promoters; ii) 50 positive and 50 negative controls of synthetically engineered sequences (*i.e.*, engineered to have multiple binding sites towards known TFs or no known binding sites, respectively)^21^; and iii) 50 positive and 50 negative controls of naturally occurring enhancers (*i.e.*, observed to exhibit high and low enhancer activity, respectively)^15^. The second library encompassed 7,500 elements in K562 cells, and consisted of: i) 6,394 putative enhancers, centered at DNase hypersensitivity peaks which did not overlap promoters; ii) 290 positive and 276 negative controls, identified by coupling CRISPRi to single cell RNA-seq measurements to identify functional enhancer-gene pairs^22^; iii) 250 negative controls derived from dinucleotide shuffling putative enhancers; iv) 50 positive and 200 negative controls of naturally occurring enhancers (*i.e.*, observed to exhibit high and low enhancer activity, respectively)^23^; and v) 24 positive and 16 negative manually selected controls in loci of interest such as α-globin, β-globin, *GATA1*, and *MYC*^24^ (**Supplementary Table 1**).

These pilot lentiMPRA libraries were used to transduce cells in triplicate, and barcodes were sequenced at the DNA and RNA levels, following a published protocol from our labs^20^. Finally, an activity score for each element was calculated as the log_2_ of the normalized count of RNA molecules from all barcodes corresponding to the element, divided by the normalized number of DNA molecules from all barcodes corresponding to the element (**Supplementary Table 2**, **Extended Data Fig. 1c**). We observed ∼50-250 median barcodes per enhancer in each replicate (**Extended Data Fig. 2b**), confirming that quantification precision was high due to the presence of a large number of independent measurements for each element. The final element activity scores, *i.e.* log_2_(RNA/DNA) ratios, were highly concordant across replicates, ranging from a Pearson correlation of 0.88-0.96 between replicate pairs (**Extended Data Fig. 2c-d)**. Averaging across the three replicates, we observed that the distribution of element activity scores was strongly divergent between most positive and negative controls (**Extended Data Fig. 2e**). An exception to this trend was observed for controls derived from CRISPRi-based screening efforts^22, 24^, which were initially chosen for testing based upon their favorable epigenetic context (*e.g.*, strong H3K27ac activity). In these cases, negative controls (*i.e.*, which displayed no significant association to a target gene) exhibited only a slightly weaker signal than positive controls (*i.e.*, which displayed significant association to a target gene), indicating that, in our reporter assays and outside their epigenetic context, they were still capable of activating transcription relative to dinucleotide shuffled negative controls (**Extended Data Fig. 2e**).

We next analyzed both lentiMPRA libraries for functional enhancers. For HepG2, we found 2,740 of the 8,960 (30.6%) measured putative enhancers to be more active than negative synthetic controls^21^ [5% false discovery rate (FDR)]. For K562, we found 3,703 of 6,315 (58.6%) measured putative enhancers to be more active than shuffled negative controls (5% FDR). Given the extensive prior work characterizing regulatory elements in the β-globin locus, and the inclusion of these sequences in our K562 library, we evaluated whether our MPRA results reproduced the findings of past work for five previously characterized CREs, termed HS1-5 (**Fig. 1a**). Consistent with prior work^25, 26^, we observed that HS2 strongly activated transcription relative to HS1 and HS3-5 (**Fig. 1a**). In summary, these pilot experiments confirmed the ability of our revised lentiMPRA protocol to measure regulatory activity with high precision and reproducibility, and recapitulated known functional features of the β-globin locus.

**Fig. 1:**
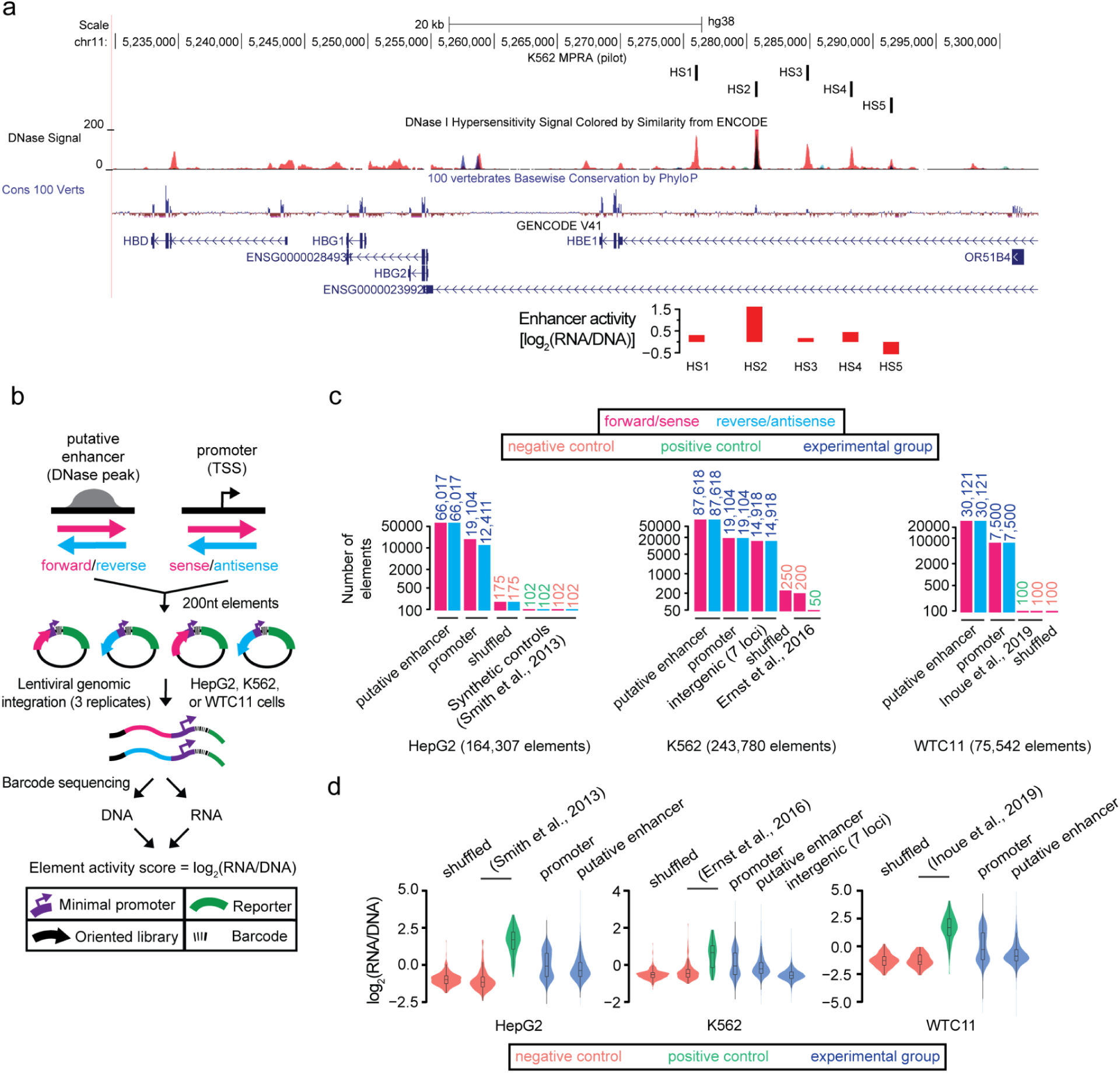
Comprehensive functional validation of CREs in three ENCODE cell types using lentiMPRA. **a,** UCSC genome browser tracks annotating, from top to bottom: i) five enhancers (HS1-HS5) of the globin locus tested in the pilot K562 MPRA library; ii) DNase hypersensitivity signal in K562 cells; iii) base conservation among 100 vertebrate species; iv) gene models in the locus shown; and v) enhancer activity scores from the pilot K562 MPRA library for each of the five enhancers tested. **b,** Schematic of lentiMPRA strategy for large-scale libraries. Thousands of CREs including putative enhancers (*i.e.*, marked by DNase peaks in the respective cell-type) and promoters [*i.e.*, centered at the transcriptional start site (TSS) of protein-coding genes] are inserted in a reporter plasmid in both orientations together with barcodes. The libraries are infected into HepG2, K562, and WTC11 cells using lentivirus, and the integrated DNA and transcribed RNA barcodes are sequenced to quantify CRE activity. **c,** Composition of the HepG2, K562, and WTC11 libraries. Thousands of putative enhancers and promoters, negative controls (dinucleotide shuffled sequences or elements lacking a signal from prior studies), and positive controls (elements with reported activity from prior studies) are included in each library^21, 23, 27^. Bars are colored according to orientation tested, with accompanying numbers indicating the number of elements tested in each category. Numbers are colored according to element type. **d,** Violin plots of element activity, measured as log_2_(RNA/DNA) ratios, for CREs, negative controls, and positive controls for each library.

### lentiMPRA CRE characterization in three cell lines

With our pilot libraries showing reproducible and robust results, we next set out to test whether our revised lentiMPRA approach could assay the majority of CREs of any given human cell type, *i.e.* over 200,000 sequences, in a single experiment. Using a similar scheme as before, we sought to test a combination of all known 19,104 protein-coding gene promoters as well as putative enhancers in both orientations (**Fig. 1b**). In HepG2 cells, we identified 66,017 putative enhancers, and tested all promoters and putative enhancers; in K562 cells, we identified 87,618 putative enhancers, and tested all promoters and putative enhancers; in WTC11 cells, due to their reduced transduction efficiency, we tested a subset of 7,500 promoters and 30,121 putative enhancers from the 83,201 identified putative enhancers (**Fig. 1c**, **Methods**). To further interrogate whether open chromatin is required for transcriptional activation, we additionally tested 14,918 heterochromatic regions in our K562 library, selected from the ±1 megabase region around the *GATA1*, *MYC*, *HBE1*, *LMO2*, *RBM38*, *HBA2*, and *BCL11A* loci, all known human disease associated and erythroid lineage genes. Collectively, incorporating dinucleotide shuffled negative controls and other positive and negative controls identified in prior studies^21, 23, 27^, we designed and tested a total of 164,307 elements in HepG2 cells, 243,780 elements in K562 cells, and 75,542 elements in WTC11 cells (**Supplementary Table 3**, **Fig. 1c**).

We observed ∼20-50 median barcodes per enhancer in each replicate among all cell types (**Extended Data Fig. 3a**). Element activity scores were also strongly concordant across replicate pairs, with Pearson correlations of about 0.94 for HepG2 cells, 0.76 for K562 cells, and 0.76 for WTC11 cells (**Supplementary Table 4**, **Extended Data Fig. 3b-d**). Averaging across the three replicates, we also observed strong agreement among element activity scores between CREs common to both our pilot and large-scale libraries (Pearson *r* = 0.94 in HepG2 cells and *r* = 0.81 in K562 cells, **Extended Data Fig. 3e**). We observed that the distribution of element activity scores was strongly divergent between positive and negative controls in each cell type, with promoters and putative enhancers spanning the range in between both extrema (**Fig. 1d**). Promoters exhibited, on average, higher activity scores and a bimodal distribution compared to putative enhancers, which exhibited a right-skewed distribution in all cell types (**Fig. 1d**). This bimodal distribution was likely caused by inactive promoters exhibiting little to no activity in the MPRA. We next analyzed all libraries to empirically measure the proportion of functional CREs among each element type and cell type. Using shuffled controls as a background set in each cell type and both orientations of measured CREs as a foreground set, we found over 50% of all promoter sequences to have regulatory activity [HepG2: 11,367 of 20,816 (54.6%); K562: 15,362 of 29,376 (52.3%); WTC11: 5,038 of 9,964 (50.6%); 5% FDR]. For enhancers, we found as many as 42% to be active [HepG2: 50,714 of 118,433 (42.8%); K562: 69,820 of 169,260 (41.3%); WTC11: 11,861 of 45,942 in (25.8%); 5% FDR].

### Properties of human promoters and their orientation dependence

Following procedures established in our prior work^19^, we utilized our substantially expanded MPRA data to examine the relative orientation dependence of promoters and enhancers. Towards this goal, we quantified the degree to which CREs exhibited observable orientation dependence. In all cell types examined, CREs cloned in the same orientation (*i.e.*, forward vs. forward or reverse vs. reverse) exhibited a correlation of ∼0.2 greater among replicate pairs than CREs cloned in the opposite orientation with respect to the reporter (*i.e.*, forward vs. reverse or reverse vs. forward) (**Fig. 2a**). When averaging among replicates in HepG2 (*i.e.*, the cell type with the greatest technical reproducibility among replicates), we detected a substantial number of CREs which exhibited greater activity in one orientation relative to the other (**Fig. 2b**). These findings suggest that the activity of CREs is largely, but not completely, independent of orientation.

**Fig. 2:**
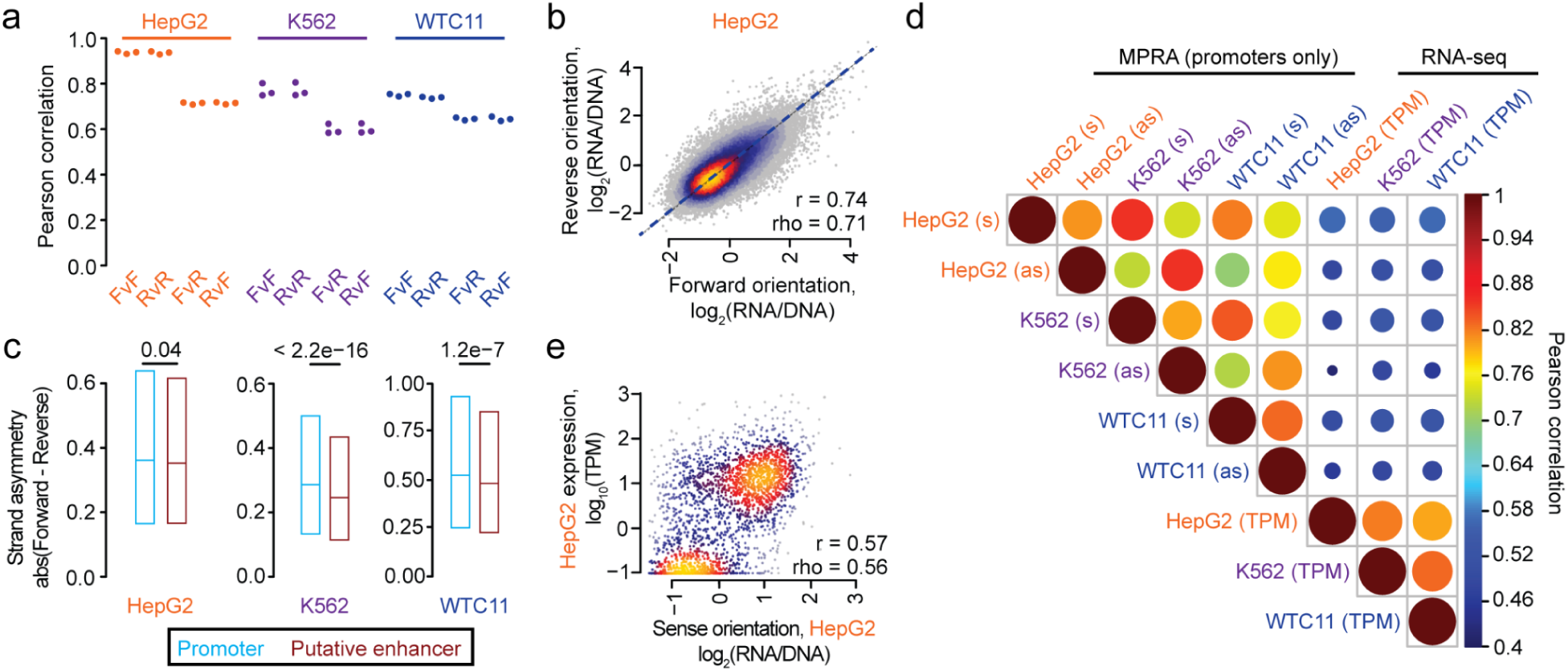
Orientation-dependent properties of human enhancers and promoters. **a,** Beeswarm plot of the Pearson correlation values corresponding to each of the three pairwise comparisons among the three replicates. The correlations are computed between observed CRE activity values for elements positioned either in the same [Forward (F) vs. F and Reverse (R) vs. R)] or opposite (F vs. R and R vs. F) orientations. **b,** Scatter plots of the average activity score for each CRE in the F vs. R orientation. Regions are colored according to the density of data from light blue (low density) to yellow (high density). Also indicated are the Pearson (r) and Spearman (rho) correlation values. **c,** Boxplots showing the distribution of strand asymmetries for promoters and putative enhancers for each cell type. Indicated is the median residual value (bar) and 25th and 75th percentiles (box). The difference between each pair of distributions was evaluated with a one-sided Wilcoxon rank-sum test. **d,** Upper triangular heatmap indicates the correlation between the sense (s) and antisense (as) orientations of promoters tested in the MPRA as well as endogenous gene expression levels measured in transcripts per million (TPM) using RNA-seq. The sizes of the circles are proportional to the Pearson correlations. **e,** Scatter plots of activity scores for sense-oriented promoters tested in the MPRA and endogenous gene expression levels for HepG2 cells. Expression levels follow a bimodal distribution.

To further compare the properties of strand asymmetry between promoters and enhancers among each element type, we analyzed strand asymmetry distributions, defined as the absolute deviation between activity scores from one orientation to the other. Consistent with prior work^28, 29^, we observed that promoters displayed greater strand asymmetry effects relative to putative enhancers in all cell types examined (**Fig. 2c**), supporting the conclusion that they contain TFBSs that promote transcription unidirectionally (or at least more unidirectionally than enhancers).

Given the enhanced orientation dependence of promoters, we sought to evaluate the relationship between orientation-specific promoter activity as measured by both the lentiMPRA and RNA-seq of endogenous genes. Across all pairs of cell types, MPRA measurements from the same orientation (*i.e.*, sense vs. sense or antisense vs. antisense) displayed greater correlation than measurements from the opposite orientation (*i.e.*, sense vs. antisense) (**Fig. 2d**). Furthermore, when comparing against endogenous expression levels, we observed: i) MPRA measurements from the matched cell type displayed nearly the same correlations as those from a different cell type; and ii) for each cell type, MPRA measurements from the sense orientation displayed greater correlation to endogenous gene expression levels than promoters tested in the antisense orientation (**Fig. 2d**). These results indicate that core promoters possess a reproducible orientation dependence and little cell-type specificity.

Despite relating to a very short promoter definition, the 200 bp core promoters centered at the transcriptional start site (TSS) and tested with lentiMPRA strongly recapitulated endogenous gene expression levels (Pearson *r* ≈ 0.55, **Fig. 2e**, **Extended Data Fig. 4a-b**). Due to the switch-like (*i.e.*, “on/off”) state of promoters, expression values fell into two modes (*i.e.*, a bimodal distribution), which slightly inflated these correlations. Removing all non-expressed genes led to a modest reduction in the correlation between MPRA measurements of promoter activity and endogenous expression levels (Pearson *r* ≈ 0.43, **Extended Data Fig. 4c-f**). Interestingly, in WTC11 cells, we found a larger cohort of transcriptionally active genes whose promoters were inactive in our MPRA (**Extended Data Fig. 4b,f**), suggestive of a shift towards a euchromatic promoter state in this cell line. Overall, motif analysis discovered a set of motifs enriched in the most highly vs. lowly expressed promoters; these were dominated by CpG-rich motifs as well as TFBSs for ETV5/7, SP1/2/4, NFYA/B/C, and THAP11/STAT5B (**Extended Data Fig. 4g**). While we did not anticipate such short 200 bp promoter fragments to reflect endogenous expression levels, collectively, these findings are consistent with prior models which showed that CpG-rich promoters are associated with elevated gene expression; and that core promoters centered at the TSS possess weak cell-type specificity, are information dense, and strongly predict gene expression levels^30^.

### Biochemical features predict regulatory activity

We next set out to train regression models that can characterize regulatory activity. Previous work by our labs trained regression models to characterize enhancer activity^15, 19^, but were limited in sample size (*i.e.*, ∼3,000 sequences) and underlying sampling bias (*e.g.*, requiring elements to have a strong H3K27ac signal). Here, we took advantage of our sizable MPRA to comprehensively test which biochemical signals from the matched cell type explain CRE activity (*i.e.*, specifically, the subset of promoters and putative enhancers). We generated lasso regression models for each cell type based upon a set of 1,506 HepG2 biochemical descriptors; 1,206 K562 descriptors; and 277 WTC11/H1-ESC descriptors (**Supplementary Table 5**). These descriptors encompass features from DNase hypersensitivity, histone ChIP-seq, and TF ChIP-seq datasets. We used these biochemical experiments to compute a feature set by extracting signal intensities from genomic regions corresponding to our elements, and then averaging signals from the samples for the identical TF or histone mark in order to reduce feature redundancy. This led to a total of 655 HepG2 features, 447 K562 features, and 122 WTC11/H1-ESC features considered by the models. We were able to predict enhancer activities with high accuracy (Pearson *r* ≈ 0.72) in all three cell types (**Fig. 3a**) using a 10-fold cross-validation approach on our data. Many of the top coefficients fit by these models correspond to DNase/ATAC-seq signal as well as ChIP-seq signal for transcriptional activators (*e.g.*, HNF1A/HNF1B/HNF4A, NFYB/NFYC, SP1, YY1, and NRF1), coactivators (*e.g.*, EP300 and KDM5A), repressors (REST, SUZ12, SIN3A, HDAC1/2), chromatin organizers often found in insulators (CTCF and RAD21), and core transcriptional machinery proteins (POLR2G and TAF1) (**Fig. 3b**). Despite these findings, model interpretation was inherently limited by the substantial degree of multicollinearity among features, as previously observed^19^. To minimize the chances that other important factors may have been overlooked by lasso regression, we identified alternative features which were highly correlated to the top-ranked features of our models (**Extended Data Fig. 5**). This analysis revealed a large suite of TFs whose activity was strongly correlated to the top features, and identified other relevant components of cohesin, including SMC3 and STAG1 (**Extended Data Fig. 5a**).

**Fig. 3:**
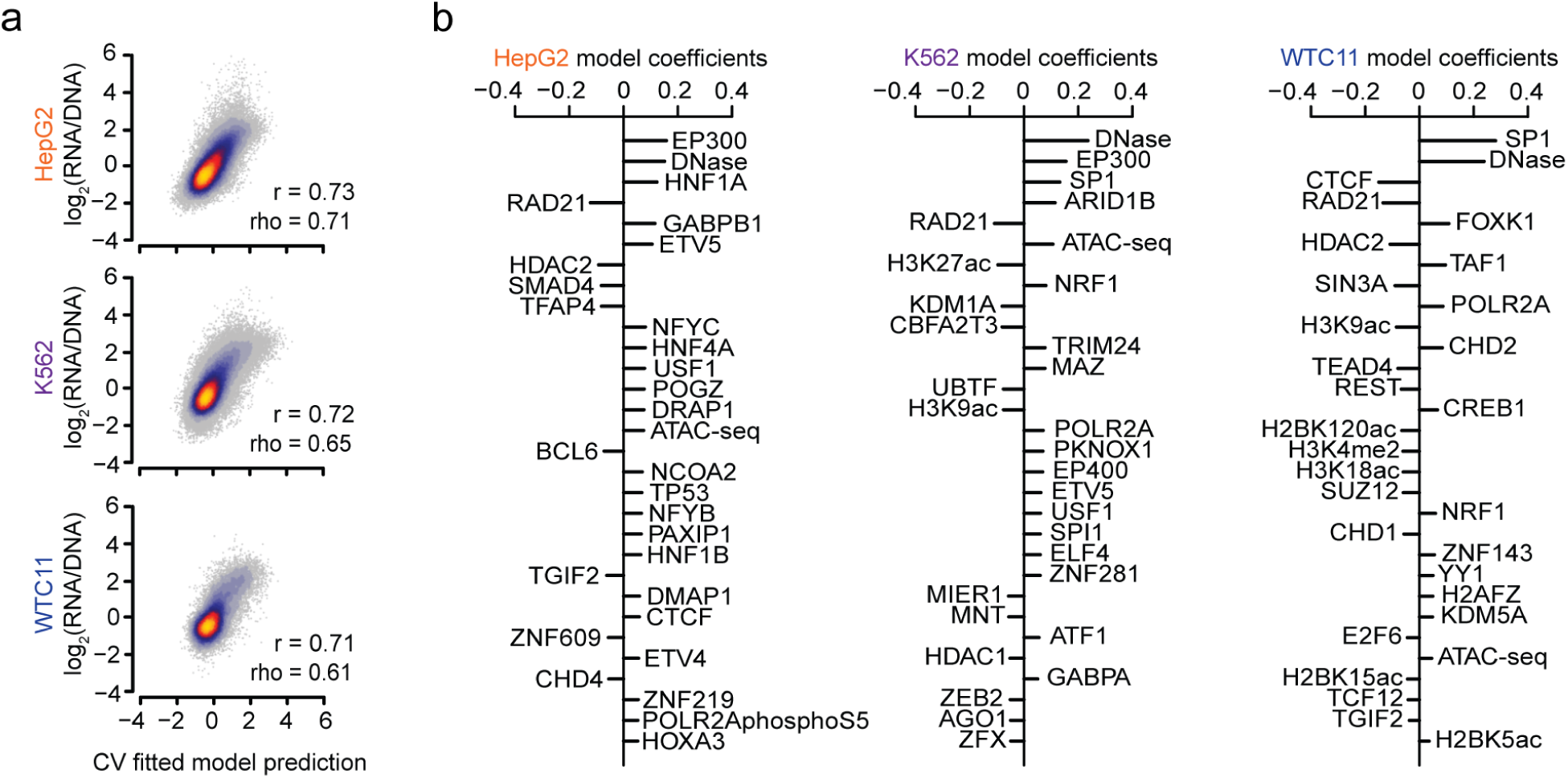
Prediction of sequence activity based on biochemical features. **a,** Scatter plot indicating relationship between model predictions and observed element activity scores for each cell type. Pearson (r) and Spearman (rho) correlation values are shown after concatenating the observations for all 10 cross-validation folds of held-out data. **b,** The top 30 coefficients derived from lasso regression models trained independently on each cell type.

### Sequence-based models predict regulatory activity with higher precision

Given the success of sequence-based models relative to biochemical models^31^, we next sought to determine the degree to which a model based on sequence features could predict the data relative to a model based on biochemical features. Towards this goal, we benchmarked the performance of two sequence-based models: i) MPRAnn, a convolutional neural network (CNN) trained on our MPRA data for each of the three cell types (**Fig. 4a**, **Extended Data Fig. 6a**); and ii) EnformerMPRA, which uses the CNN-transformer architecture Enformer^32^ to generate a feature set of 5,313 biochemical predictions and then fit a lasso regression to the MPRA data using these features. Comparing the performance of MPRAnn and EnformerMPRA to our biochemical lasso regression model on the identical 10 folds of held-out data, we observed that MPRAnn outperformed the biochemical model, and EnformerMPRA outperformed MPRAnn in each of the three cell types (**Fig. 4b**). Combining the folds of data, our best model EnformerMPRA achieved a performance (Pearson *r* ≈ 0.84, **Fig. 4c**) comparable to the technical noise of the assay itself (*i.e.*, the replication of replicates, **Extended Data Fig. 3b-d**). Despite its superior performance, the multicollinearity of the feature set, coupled to the large number of features, obfuscated a mechanistic understanding of which TFBSs drive EnformerMPRA’s predictions. We therefore used MPRAnn, which can be easily deployed to generate *in silico* mutagenesis (ISM) predictions while circumventing the need to compute an extensive feature set. Generating ISM predictions on the full set of MPRA data, we used TF-MoDISco-lite^33^ to identify motifs at variants with a large predicted effect size. This strategy identified a large suite of housekeeping TFs predicted to activate transcription in all cell types, including JUN, NRF1, CEBPA/D/G, USF1, THAP11, and ELF1; additionally, we discovered a motif for REST, a known transcriptional repressor^34^, predicted to inhibit transcription in HepG2 and K562 cells (**Extended Data Fig. 6b**). The top three TFBSs most frequently associated with transcriptional activation among all cell types corresponded to KLF5/15, NFYA/NFYC, and FOXI1/FOXJ2; in contrast, the top cell-type-specific TFBS corresponded to HNF4A/G in HepG2 cells, GATA2/3 in K562 cells, and POU5F1::SOX2 in WTC11 cells (**Fig. 4d**).

**Fig. 4:**
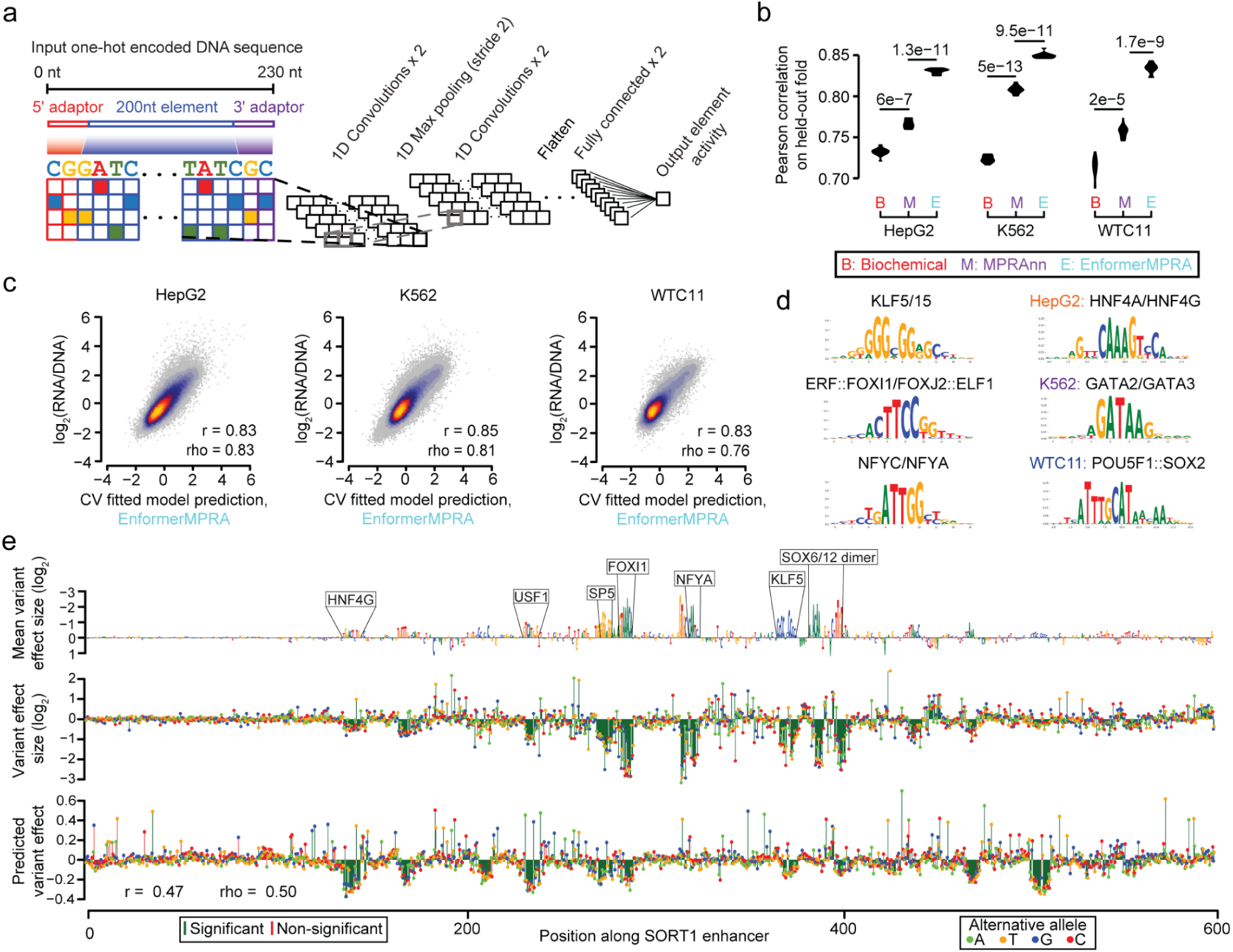
Sequence-based models predict regulatory element activity. **a,** MPRAnn is a deep convolutional neural network architecture trained to predict CRE activity from an input sequence of the tested element. **b,** Violin plots showing the performances of the trained MPRAnn and EnformerMPRA models on each of the ten cross-validation folds of held-out data, relative to the corresponding performances from our biochemical lasso regression models. An improvement relative to another model was evaluated with a one-sided, paired t-test. **c,** Scatter plot indicating relationship between EnformerMPRA model predictions and observed element activity scores for each cell type. Also indicated are the Pearson (r) and Spearman (rho) correlation values after concatenating the observations for all ten folds of held-out data. **d,** Set of enriched motifs discovered by TF-MoDISco-lite^33^; shown on the left are the top three motifs detected across multiple cell types (*i.e.*, bound by housekeeping TFs) and on the right the top motif detected for each cell type. See also **Extended Data Fig. 6b** for a comprehensive list of detected motifs. **e,** Saturation mutagenesis data from the *SORT1* enhancer^35^. Shown in the top row is the reference sequence scaled to the mean effect size among all alternative mutations, annotated by seven out of thirteen significant TFBSs that match known motifs^36^. Measured effect sizes of individual variants are displayed in the second row. The bottom row shows MPRAnn predictions as well as corresponding Pearson (r) and Spearman (rho) correlation values to the observed data.

Next, we sought to validate the accuracy of our ISM scores on promoters (*F9*, *LDLR*, *PKLR*) and an enhancer (*SORT1*) for which we had previously applied MPRA saturation mutagenesis to measure the effect sizes of all possible variant effects of these CREs in HepG2 (*F9*, *LDLR*, *SORT1*) or K562 (*PKLR*) cells^35^. Comparing MPRAnn predictions for the *SORT1* enhancer to MPRA data revealed that most of the relevant TFBSs (*e.g.*, HNF4G, USF1, SP5, FOXI1, and KLF5) could be detected, although the predicted effect sizes were exaggerated for other TFs (**Fig. 4e**). Collectively, we observed a correlation of 0.47 for *SORT1*, 0.58 for *PKLR*, 0.61 for *LDLR*, and 0.59 for *F9* between model predictions and observed data (**Extended Data Fig. 6c-f**), confirming that MPRAnn, despite being trained on CRE activity, could partially model the regulatory effects of individual genetic variants.

### lentiMPRA detects factors involved in cell-type-specific activity

While our large-scale MPRAs focused upon element activity within each of several cell types, they did not directly evaluate the cell-type-specific activity of each element. To address this limitation, we designed a lentiMPRA library to test a common set of elements in each of the three cell types. This joint library consisted of ∼19,000 putative enhancers from each of the three cell lines, sampled uniformly from prior large-scale MPRA experiments to span a wide range of activity; a subset of promoters exhibiting high expression variance as well as a wide range of average expression among cell types from our prior large-scale MPRA experiments; dinucleotide shuffled controls; and a set of positive and negative controls using synthetic elements engineered to exhibit activity in HepG2 cells^21^, or natural elements with evidence to exhibit K562-specific activity^23^ (**Supplementary Table 6**, **Fig. 5a**, **Methods**). Elements were largely tested in a single orientation (*i.e.*, sense orientation for promoters and forward orientation for putative enhancers).

**Fig. 5:**
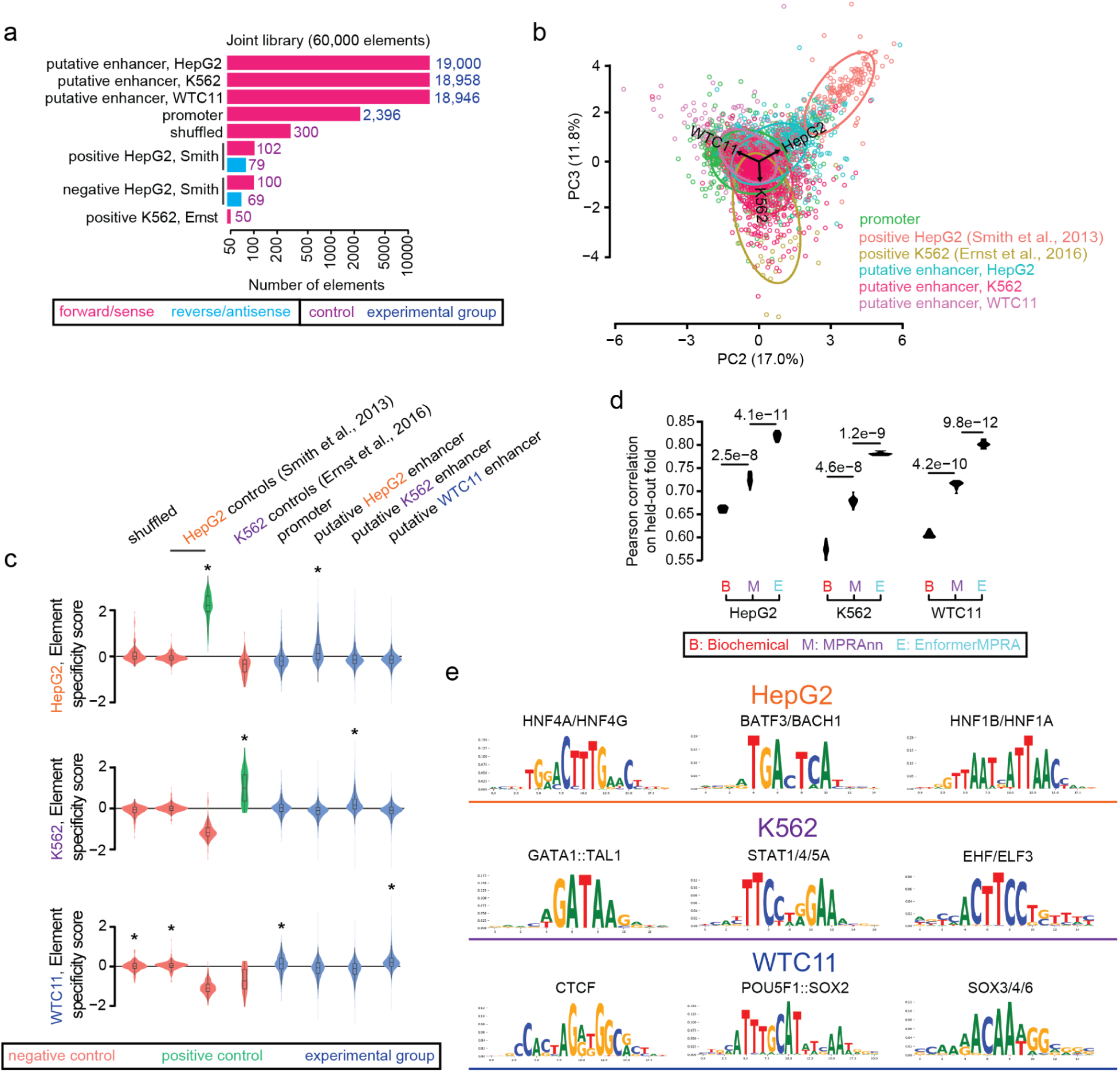
Assessment of CRE cell-type-specific activity. **a,** Composition of the joint library tested in HepG2, K562, and WTC11 cells. A similar proportion of putative enhancers sampled from each cell type were tested as well as a smaller number of promoters, negative controls (dinucleotide shuffled sequences or elements lacking a signal from prior studies), and positive controls (elements with reported activity from prior studies) are included in each library^21, 23^. **b,** PCA biplot indicating the second (PC2) and third (PC3) principal components, with a random sample of up to 1,000 data points sampled from each element category tested. The loading vectors (*i.e.*, corresponding to each cell type) as well as ellipses fitting the regions of highest density for each element category are also shown. **c,** Violin plots showing the distribution of element specificity scores for each element category, alongside information about which distributions shows a median significantly greater than zero (one-sided Wilcoxon signed-rank test, *p < 0.05). **d,** Performance of trained MPRAnn and EnformerMPRA models on each of ten cross-validation folds of held-out data, relative to the corresponding performance of lasso regression models trained on biochemical features. An improvement relative to another model was evaluated with a two-sided, paired t-test. **e,** Set of top three motifs enriched in each cell type discovered by TF-MoDISco-lite^33^. See **Extended Data Fig. 11** for a complete list of detected motifs.

We observed ∼20-50 median barcodes per enhancer in each replicate among all cell types (**Extended Data Fig. 7a**). Element activity scores were also strongly concordant across replicate pairs, with Pearson correlations of 0.98 for HepG2 cells, 0.97 for K562 cells, and 0.96 for WTC11 cells (**Supplementary Table 7**, **Extended Data Fig. 7b-d**). Averaging across the three replicates, we also observed strong agreement among element activity scores between CREs common to both our joint and large-scale libraries (Pearson *r* = 0.90 in HepG2 cells, *r* = 0.88 in K562 cells, and *r* = 0.83 in WTC11 cells **Extended Data Fig. 7e**). We observed the distribution of element activity scores to be strongly divergent and modestly cell-type-specific between positive and negative controls in each cell type (**Extended Data Fig. 8**). Although promoters and putative enhancers displayed significant activity in all cell types, the distribution of activities for putative enhancers derived from the matched cell type was greater than those from unmatched cell types (**Extended Data Fig. 8**). To examine cell-type specificity in more detail, we evaluated the behavior of each element category (*i.e.*, promoters or putative enhancers from each cell type) in each pair of cell types. Promoters exhibited the strongest correlation among cell-type pairs (mean Pearson *r* = 0.82); in contrast, putative enhancers displayed weaker correlations when comparing the activity scores from the cell type from which the enhancer was derived to those from a different cell type (Pearson *r* = 0.32–0.51 for HepG2 enhancers, *r* = 0.51–0.65 for K562 enhancers, and *r* = 0.64–0.65 for WTC11 enhancers, **Extended Data Fig. 9**). Collectively, our results show that promoters are less cell-type specific, likely due to the presence of TFBSs for housekeeping TFs which can universally activate them. In contrast, enhancers show stronger cell-type specificity, in line with their presumed cell-type-specific functions^37^.

We next sought to more deeply interrogate the cell-type-specific activity of each element. Towards this goal, we performed a principal components analysis (PCA) using our matrix of element activity scores in three cell types, and removed the dominant PC, which represented the “universal” signal of element activity among cell types. An analysis of PC2 and PC3 revealed that promoters have a slight bias towards expression in WTC11 cells, and both controls as well as putative enhancers have a stronger bias towards greater expression in the cell type from which they were derived (**Fig. 5b**). We next computed an element specificity score, which measures each element’s deviation from its mean activity across cell types. These scores recapitulated the expected patterns of enrichment or depletion of element activity for different element categories, with HepG2 and K562 controls showing strong relative activity in their respective cell types; putative enhancers showing strong relative activity in their respective cell types; and promoters and negative controls showing modestly stronger activity in WTC11 cells relative to others (**Fig. 5c**, **Extended Data Fig. 10**).

We next benchmarked the performance of biochemical and sequence-based models in predicting our element specificity scores from this lentiMPRA library. Consistent with prior results, a multi-task version of MPRAnn outperformed the biochemical model, while EnformerMPRA outperformed both MPRAnn and the biochemical model for each of the three cell types (Pearson *r* ≈ 0.81 for EnformerMPRA; **Fig. 5d**). Using TF-MoDISco-lite^33^, we identified cell-type-specific motifs learned by our multi-task MPRAnn, detecting 17 motifs to be associated with cell-type-specific activation in HepG2 cells, and 9 motifs each in both K562 or WTC11 cells (**Extended Data Fig. 11**). The top three ranked cell-type-specific TFBSs from each cell type include HNF4A/G, BATF3/BACH1, and HNF1A/B in HepG2 cells; GATA1::TAL1, STAT1/4/5A, and EHF/ELF3 in K562 cells; and CTCF, POU5F1::SOX2, and SOX3/4/6 in WTC11 cells (**Fig. 5e**). An examination of the gene expression levels of these TFs in our three cell types revealed that *HNF4A/G*, *HNF1A/B*, *GATA1*, *TAL1*, *STAT5A*, *SOX2*, and *SOX3* exhibited strong cell-type-specific expression in the expected cell types; additionally, CTCF showed modestly enriched expression in WTC11 cells.

## DISCUSSION

MPRA is a high-throughput technology that enables the multiplex testing of thousands of sequences and variants for their regulatory activity. Here, by modifying our lentiMPRA protocols, we were able to test over 200,000 sequences in a single experiment, a number that is in line with the majority of CREs detected via descriptive biochemical gene regulatory element detection methods for a given cell type. By testing all protein-coding promoters, we find significant promoter activity bias in terms of strand orientation. We also observed that a 200 bp core promoter was sufficient to drive regulatory activity that was in line with endogenous gene expression levels. Testing both biochemical and sequence-based models, we found the sequence-based models to be superior in predicting element activity. By further testing 60,000 sequences in all three cell types, we found promoters to be more ubiquitous in their activity compared to enhancers. We also identified TFBSs that appear to convey cell-type-specific activity on enhancers.

While other large-scale MPRA datasets are available for other cell lines^11–14^, they are primarily tested via episomal STARR-seq, require a very large number of cells, only provide an episomal readout, and tend to use a strong promoter to increase the ability to detect activity^12^. In contrast, our lentiMPRAs provide nearly comprehensive functional datasets for these three cell types tested with an ‘in genome’ readout. The ability to systematically test the majority of CREs in an unbiased manner for a given cell type allowed us to identify predictive biochemical and sequence-based features for each cell type with high confidence.

Our large-scale lentiMPRAs allowed us to comprehensively test all known 19,104 protein-coding gene promoters in both orientations in two cell lines (HepG2 and K562) and 7,500 in a third cell line (WTC11). In addition to observing significant promoter activity bias in terms of strand orientation in line with previous work^28, 29^, this allowed us to extensively characterize the sequence-based information needed to generate these ‘on/off’ switches. We found that 200 bp blocks centered at the TSS of protein-coding genes have sufficient sequence data to provide this switch and are sufficient to drive expression in a similar manner to their associated gene. Dissection of the sequence content in these active core promoters versus inactive cores found them to be enriched for CpG rich motifs that are known to have ubiquitous function^30^ (**Extended Data Fig. 4g**). They also include the SP (SP1/2/4) family of TFs that are known to interact with the transcription initiation complex, epigenetic regulatory and many other TFs to provide ubiquitous promoter activity^38^ and the ETV (ETV5/7) family which is known to be enriched in ubiquitously expressed promoters^39^. We also observed an enrichment for the NF-Y (NFYA/B/C) family that is known to interact with the CCAAT box, usually located –89 bp and –50 bp from the TSS in TATA-containing and TATA-less eukaryotic promoters, respectively^40^. Of note, our lentiMPRA design tested promoters along with a minimal promoter that is 32 bp long which could affect promoter activity. However, this approach enabled us to test *en masse* hundreds of thousands of enhancers along with thousands of promoters and compare them in the same assay. Our results were similar to previous reports^28, 29^, showing orientation biases for promoters, and identified an enrichment for motifs that are known to provide ubiquitous promoter expression, further supporting that the addition of these 32 bp to our assayed promoters likely did not affect our findings.

Our large-scale and unbiased testing of hundreds of thousands of sequences for enhancer activity in three different cell lines in two different orientations (along with testing 60,000 sequences in all three cell lines) led to the following key findings: 1) enhancer activity is largely, but not completely, independent of orientation, in line with part of the original definition of enhancers^41^ and repressors. 2) enhancers have more inherent cell-type specificity than promoters; 3) cell-type specificity is driven by a small number of cell-type-specific TFBSs in each cell line. Of note, due to oligonucleotide synthesis constraints, while we were able to test hundreds of thousands of sequences in a cost-effective way, our assayed sequences were only 200 bp long, which only allowed us to test the central sequences around their DNase peak. This likely led to the enrichment observed for active sequences around open chromatin and could also limit our ability to detect cell-type-specific motifs. Furthermore, previous work by our labs^19^ showed that sequence length can have a major effect on regulatory activity in MPRA.

In line with previous work^15, 19, 42^, we show that sequence-based models provide superior ability to predict functional sequences from MPRA. Due to the large-scale nature of these lentiMPRAs, our results however provide stronger support for this observation. EnformerMPRA could not detect the features driving these predictions, due to the multicollinearity of the feature set. However, MPRAnn, while achieving slightly lower prediction scores, allowed us to tease out many motifs that are important for these predictions. These include universal TFBSs along with cell-type-specific motifs, which in combination allow for strong regulatory activity. Interestingly, one of the main TFs that were found to be enriched in both our promoters and enhancers in all three cell types are the “stripe” TFs KLF5/15. “Stripe” TFs are thought to provide co-accessibility and increase residence time for other transcription-associated factors in promoters and enhancers^43^ and were also found to be enriched in active regulatory elements in a recent large-scale lentiMPRA carried out in primary developing neurons and cerebral organoids^44^, in line with their generalizable function.

In summary, our work provides a nearly comprehensive catalog of functional regulatory elements in three established cell lines that are widely used in biological research: HepG2, K562, and iPSCs. These datasets will be extremely beneficial for developing machine learning-based tools that predict regulatory activity. They will also improve our understanding of the gene regulatory code and decode how variants within these sequences can lead to specific phenotypes. Our data could also be beneficial for future large-scale MPRAs that mutate specific motifs in known regulatory elements or use synthetic motifs to decode regulatory logic. In addition, it could be utilized to build tissue-specific regulatory elements that are designed to provide regulatory activity at various levels, similar to dimmer switches, or cell-type-specific regulatory elements that could be utilized for therapeutic delivery.

## METHODS

### Design of Agilent oligonucleotide library

#### HepG2 pilot library

For the HepG2 pilot library, we collected two replicates of DNase I hypersensitivity data derived from HepG2 cells (ENCODE narrowPeak BED files: ENCFF505SRS and ENCFF268DTI, hg19 human genome build)^45^. For each replicate, we collapsed overlapping peaks using bedtools merge (parameters “-o collapse -c 2,3,7”). Then, we identified peaks that intersected between the two replicates, merged these peaks, and removed the subset of merged peaks that overlap promoters (defined as regions ±2,500nt around any annotated transcriptional start site). The resulting distribution of peak sizes was such that 97% of peaks were ≤ 200 bp in length. We therefore centered the designed oligos at each merged DNase peak, consistent with the known region of maximal regulatory activity^23^, and added ±100 bp to either side. This procedure resulted in a set of 66,017 CREs. For this pilot library, we sought to evaluate CREs which overlapped a wide range of putative TF binding sites. We therefore intersected these putative enhancers with wgEncodeRegTfbsClusteredWithCellsV3.bed.gz^46^ in order to count the number of putative HepG2 TF binding sites intersecting these CREs. We uniformly and randomly sampled ∼1,834 CREs with 0-1, 1-5, 5-10, 10-20, and >20 TF binding sites for a total of 9,172 elements. Including 50 positive and 50 negative controls from each of two prior studies^15, 21^ resulted in a total of 9,372 elements. These 171 bp controls from prior work were linked downstream of a 29nt random sequence GGTGCTCGATTTAATTTCGCCGACGTGAT to match the 200 bp sequence length of CREs. For the final oligo library, each element was linked to the 5′ adaptor AGGACCGGATCAACT and 3′ adaptor CATTGCGTGAACCGA, designing two 230 bp oligos per element to minimize the impact of oligo synthesis errors.

#### K562 pilot library

An analogous procedure was followed for the K562 pilot library as in the “HepG2 pilot library” section, with the following modifications: i) ENCODE narrowPeak BED files ENCFF027HKR and ENCFF154JCN (hg38 human genome build)^45^ were used; ii) merging these peaks resulted in 34,367 putative enhancers; iii) after intersecting the peaks with K562 TF binding sites, we sampled ∼1,278 enhancers from each TF binding site bin to test a total of 6,394 CREs; iii) additional negative controls were chosen by dinucleotide shuffling the enhancer set; iv) positive and negative controls were chosen from CRISPRi screens^22, 24^, a prior MPRA^23^, and select loci of interest such as α-globin and β-globin; v) a total of 7,500 elements were tested; and vi) controls were already 200 bp in length, requiring no addition of a random sequence.

#### HepG2 large-scale library

Following the procedures outlined for the “HepG2 pilot library” section, we tested all 66,017 previously identified CREs in both orientations. For human protein-coding gene promoters, we extracted the average signal across cell types in Transcripts Per Million (TPM) for each CAGE peak listed in hg19.cage_peak_phase1and2combined_tpm_ann.osc.txt.gz from the FANTOM5 consortium^47, 48^. The first exons of all protein-coding gene transcripts were collected from Ensembl v83 (hg38 genome build)^49^, transformed into hg19 coordinates using liftOver^50^, and then intersected with the CAGE peaks to identify a single promoter per gene corresponding to the promoter with the maximal average TPM. To select the final 200 bp oligo for testing, we identified the center of the promoter DNase peak based upon the HepG2 DNase peaks merged across replicates (described in the “HepG2 pilot library” section). In the scenario in which no DNase peak overlapped the promoter, we centered upon the midpoint of the CAGE peak. In the scenario in which neither a DNase nor CAGE peak existed, we centered upon the TSS defined by the Ensembl annotation. This resulted in a total of 19,104 protein coding gene promoters, of which 6,181 were centered on a DNase peak, 9,735 were centered on a CAGE peak, and 3,188 were centered on a Ensembl TSS definition. The oligo tested included the ±100 bp window around this central position in the sense orientation with respect to the gene. A random subset of 12,411 promoters were also tested in the antisense orientation. We tested 102 positive and 102 negative controls from a prior study^21^ as well as 175 dinucleotide shuffled negative controls in both orientations. This resulted in a library consisting of 164,307 elements, for which we ordered one 230 bp oligo per element.

#### K562 large-scale library

To acquire a set of DNase peaks for testing, we used the “optimal peak” calls derived from processing ENCODE experiment ID: ENCSR000EOY through the ENCODE DCC Irreproducible Discovery Rate (IDR) pipeline, available at https://github.com/ENCODE-DCC/atac-seq-pipeline (generously provided to us by A. Kundaje). Removing DNase peaks overlapping human promoters resulted in 87,618 putative enhancers tested in both orientations. The promoters tested were identical to those described in the “HepG2 large-scale library”, except that it included all 19,104 promoters tested in both orientations. We tested 50 positive and 200 negative controls from a prior MPRA study^23^ as well as 250 dinucleotide shuffled negative controls. Finally, 14,918 tiles not overlapping DNase peaks, and subsampled from the ±1 Megabase region around the following 7 genetic loci: *GATA1*, *MYC*, *HBE1*, *LMO2*, *RBM38*, *HBA2*, and *BCL11A*, were chosen using our representative subset selection approach (*i.e.*, described below) and tested in both orientations. This resulted in a library consisting of 243,780 elements, for which we ordered one 230 bp oligo per element.

#### WTC11 large-scale library

To acquire a set of DNase peaks for testing, we used the peak calls derived from applying the hotspot2 pipeline (https://github.com/Altius/hotspot2) at FDR = 0.05 to ENCODE experiment ID: ENCSR785ZUI (generously provided to us by R.Sandstrom)^51^. This resulted in two independent replicates, which were merged into a unified set using the procedures described in the “HepG2 pilot library” section. Removing DNase peaks overlapping human promoters resulted in 83,201 putative enhancers. Together with the 19,104 promoters described in the “HepG2 large-scale library”, these elements were subsampled to select 30,121 putative enhancers and 7,500 promoters using our representative subset selection approach described below, and tested in both orientations. We also tested 100 positive and 100 negative controls from a prior study^27^ as well as 100 dinucleotide shuffled negative controls. This resulted in a library consisting of 75,542 CREs, for which we ordered one 230 bp oligo per element.

#### Joint library tested in HepG2, K562, and WTC11 cells

Given the measured putative enhancers from the forward orientations in each of the HepG2, K562, and WTC11 large-scale libraries, we binned each set of CREs into ten equally sized bins spanning the range of measured log_2_(RNA/DNA) values in the selected cell type. We randomly sampled an approximately equal number from each bin, resulting in 19,000 HepG2, 18,958 K562, and 18,946 WTC11 putative enhancers. A similar procedure was followed with sense-oriented promoters, except that the ten bins were established based upon the mean log_2_(RNA/DNA) across all three cell types (*i.e.*, instead of performing the procedure independently in each cell type as before), and the top 1,000 promoters exhibiting the greatest variance across three cell lines were also selected for testing. This resulted in the selection of 2,396 out of 19,104 promoters. We also tested 181 positive and 169 negative HepG2 controls from a prior study^21^, 50 positive K562 controls from a prior study^23^, and 300 dinucleotide shuffled negative controls. This resulted in a library consisting of 60,000 CREs, for which we ordered one 230 bp oligo per element.

#### Representative subset selection

Given the limited number of testable elements in the large-scale K562 and WTC11 libraries, we designed a subset selection procedure to more optimally sample a non-redundant subset of elements associated with diverse biochemical features. For K562 cells, we used ground sets of non-overlapping 200 bp windows uniformly covering each of 7 genetic loci; for WTC11 cells, we used ground sets of 83,201 putative enhancers and 19,104 promoters. To perform representative subset selection with these ground sets, we utilized an objective function called facility location. This submodular set function can be optimized using a greedy algorithm, and yields a subset of elements that covers the epigenetic diversity of the ground set^52^. The facility location function is given as:

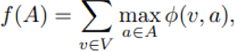

where *V* is the ground set, *A* is a subset of *V* with *k* elements and φ is a nonnegative similarity function. Optimization of the facility function was performed using python package *apricot* (https://github.com/jmschrei/apricot/). For this study, we set *k* = 2,231 for each of the 7 loci in K562 cells, *k* = 30,121 for WTC11 putative enhancers, and *k* = 7,500 for promoters in WTC11. From the 7 loci, we then filtered out the tiles that overlapped DNase peaks as they had already been tested, and then subsampled to ∼2,131 tiles per locus to retrieve 14,918 tiles among all loci.

To assess the pairwise similarity of each element, we utilized hundreds of ENCODE histone and TF ChIP-seq experiments derived from K562 and WTC11/H1-ESCs (**Supplementary Table 5**). For each 200 bp tile in the ground set, we computed the mean signal for each ChIP-seq dataset, resulting in a vector of biochemical measurements for each 200 bp tile. We used the Pearson correlation coefficient as a similarity function given these ChIP-seq features.

### Generation of MPRA libraries

The MPRA libraries were generated as previously described^20^. In brief, the Agilent oligo pool was amplified by 5-cycle PCR using forward primer (pLSmP-enh-f, **Supplementary Table 8**) and reverse primer (minP-enh-r, **Supplementary Table 8**) that adds a minimal promoter and spacer sequences downstream of the CRE. The amplified fragments were purified with 0.8x AMPure XP (Beckman coulter), and amplified for 15 additional cycles using the forward primer (pLSmP-enh-f) and reverse primer (pLSmP-bc-primer-r, **Supplementary Table 8**) to add 15 bp random sequence that serves as a barcode. The amplified fragments were then inserted into *Sbf*I/*Age*I site of the pLS-SceI vector (Addgene, 137725) using NEBuilder HiFi DNA Assembly mix (NEB), followed by transformation into 10-beta competent cells (NEB, C3020) using the Gemini X2 machine (BTX). Colonies were allowed to grow up overnight on Carbenicillin plates and midiprepped (Qiagen, 12945). For HepG2 and K562 pilot libraries, we collected approximately 1 million and 1.3 million colonies, so that on average 50 and 100 barcodes were associated with each CRS, respectively. For HepG2, K562 and WTC11 large-scale libraries, we collected approximately 8M, 12M, and 3M colonies aiming to associate approximately 50, 50, and 40 barcodes per CRS, respectively. For the joint library, we collected approximately 3.3M colonies aiming to associate approximately 55 barcodes per CRS. To determine the sequences of the random barcodes and their association to each CRS, the CRS-mP-barcodes fragment was amplified from each plasmid library using primers that contain flowcell adapters (P7-pLSmP-ass-gfp and P5-pLSmP-ass-i17, **Supplementary Table 8**). The fragment was then sequenced with a NextSeq mid-output 300 cycle kit using custom primers (Read 1, pLSmP-ass-seq-R1; Index read, pLSmP-ass-seq-ind1; Read 2, pLSmP-ass-seq-R2, **Supplementary Table 8**).

### Cell culture, lentivirus packaging, and titration

HEK293T (ATCC, CRL-3216), HepG2 (ATCC, HB-8065) and K562 (ATCC, CCL-243) cell culture were performed as previously described^15^. WTC11 human iPSCs (RRID:CVCL_Y803) were cultured in mTeSR plus medium (Stemcell technologies, Catalog # 100-0276) and passaged using ReLeSR (Stemcell technologies, Catalog # 100-0484), according to the manufacturer’s instructions. WTC11 cells were used for the MPRA experiments at passage 43-49. Lentivirus packaging and titration were performed as previously described with modifications^20^. Briefly, 50,000 cells/cm^2^ HEK293T cells were seeded in T175 flasks and cultured for 48 hours. The cells were co-transfected with 7.5 μg/flask of plasmid libraries, 2.5 μg/flask of pMD2.G (Addgene 12259), and 5 μg/flask of psPAX2 (Addgene 12260) using EndoFectin Lenti transfection reagent (GeneCopoeia) according to the manufacturer’s instructions. After 8 hours, cell culture media was refreshed and ViralBoost reagent (Alstem) was added. The transfected cells were cultured for 2 days and lentivirus were harvested and concentrated using the Lenti-X concentrator (Takara) according to the manufacturer’s protocol. To measure DNA titer for the lentiviral libraries in HepG2, K562, or WTC11, cells were seeded at 1×10^5^ cells/well in 24-well plates and incubated for 24 hours. Serial volume (0, 2, 4, 8, 16, 32 μL) of the lentivirus was added along with polybrene at the final concentration of 8 μg/ml. The infected cells were cultured for three days and then washed with PBS three times. Genomic DNA was extracted using the Wizard SV genomic DNA purification kit (Promega). Multiplicity of infection (MOI) was measured as relative amount of viral DNA (WPRE region, forward; 5′-TACGCTGCTTTAATGCCTTTG-3′, reverse; 5′-GGGCCACAACTCCTCATAAAG-3′) over that of genomic DNA (intronic region of LIPC gene, forward; 5′-TCCTCCGGAGTTATTCTTGGCA-3′, reverse; 5′-CCCCCCATCTGATCTGTTTCAC-3′) by qPCR using SsoFast EvaGreen Supermix (BioRad), according to the manufacturer’s protocol.

### Lentiviral infections and DNA/RNA barcode sequencing

For the HepG2 and K562 pilot libraries, 2.4M HepG2 or 10M K562 cells per replicate were seeded in 10cm dishes or T75 flasks, respectively, and incubated for 24 hours. The HepG2 and K562 cells were infected with the lentiviral libraries along with 8 μg/ml polybrene, with an estimated MOI of 50 or 10, respectively. For the large-scale HepG2 library, 15M HepG2 cells per replicate were seeded in three 15cm dishes (5M per dish), incubated for 24 hours, and infected with the library along with 8 μg/ml polybrene, with an estimated MOI of 50. For the large-scale K562 library, 85M K562 cells per replicate were seeded in three T225 flasks (28.3M per flask), incubated for 24 hours, and infected with the library along with 8 μg/ml polybrene, with an estimated MOI of 10. For the large-scale WTC11 library, 38.4M WTC11 cells per replicate were seeded in four 10cm dishes (9.6M per dish), incubated for 24 hours, and infected with the library along with 8 μg/ml polybrene, with an estimated MOI of 10. For the joint library, 5M HepG2, 28M K562, and 38.4M WTC11 cells were infected with the estimated MOI of 50, 10, and 10, respectively. For each experiment, three independent infections were performed to obtain three biological replicates. After three days of culture, genomic DNA and total RNA were extracted from the infected cells using AllPrep DNA/RNA mini kit (Qiagen), and sequencing library preparations were performed as previously described^20^. The libraries were then sequenced with a NextSeq high-output 75 cycle kit using custom primers (Read 1, pLSmP-5bc-seq-R1; Index1 (UMI read), pLSmP-UMI-seq; Index2, pLSmP-5bc-seq-R2; Read 2, pLSmP-bc-seq; **Supplementary Table 8**)^20^.

### MPRA processing pipeline

#### Associating barcodes to designed elements

For each of the barcode association libraries, we generated Fastq files with bcl2fastq v2.20 (parameters “--no-lane-splitting --create-fastq-for-index-reads --use-bases-mask Y*,I*,I*,Y*”), splitting the sequencing data into paired-end index files delineating the barcodes (*i.e.*, I1 and I2) and paired-end read files delineating the corresponding element linked to the barcode (*i.e.*, R1 and R2). These files were used to associated barcodes to elements using the *association* utility of MPRAflow 1.0^20^ (run as: nextflow run association.nf --fastq-insert “R1.fastq.gz” --fastq-insertPE “R2.fastq.gz” --fastq-bc “I1.fastq.gz” --fastq-bcPE “I2.fastq.gz” --aligner “bt2_strand” --design “designed_sequences.fa”). Here, “designed_sequences.fa” was a FASTA file incorporating all of the element sequences that had been ordered from the corresponding Agilent library, and ‘bt2_strand’ was used to map elements in a strand-aware fashion to accommodate the existence of elements tested in both orientations. The final output of this utility was a “filtered_coords_to_barcodes.pickle” file mapping barcodes to elements.

#### Replicates, normalization, and RNA/DNA activity scores

For each of the indexed DNA and RNA libraries, we demultiplexed the sequencing run and generated Fastq files with bcl2fastq v2.20 (parameters “--barcode-mismatches 2 --sample-sheet SampleSheet.csv --use-bases-mask Y*,Y*,I*,Y* --no-lane-splitting --minimum-trimmed-read-length 0 --mask-short-adapter-reads 0”), where “SampleSheet.csv” cataloged the correspondence between the index sequence and DNA or RNA replicate sample of origin. In several instances, the “--barcode-mismatches 2” resulted in an index assignment clash, requiring us to instead use “--barcode-mismatches 1”. These commands split the sequencing data into paired-end read files delineating the barcodes (*i.e.*, R1 and R3) and a file indicating the unique molecular index (UMI) (*i.e.*, R2) for each DNA or RNA replicate sample. We compiled a table of these files to indicate the 3 RNA and 3 DNA files for each of the three replicates in the file “experiment.csv”. Finally, we used the *count* utility of MPRAflow 1.0^20^ (run as: nextflow run count.nf --e “experiment.csv” --design “designed_sequences.fa” --association “filtered_coords_to_barcodes.pickle”) to compute activity scores for each element and replicate as log_2_(RNA/DNA). Elements with which were measured with fewer than 10 independent barcodes were removed to reduce the impact of measurement noise in downstream analysis. To combine the data from all three replicates, the distribution of activity values was normalized to the median activity value within each replicate, and then the activity values were averaged across the three replicates.

### Regression modeling

#### Biochemical model features

We extracted all TF ChIP-seq, histone ChIP-seq, DNase-seq, and ATAC-seq bigWig files available for HepG2, K562, and WTC11 cells for the hg38 human genome assembly under “released” ENCODE status^46^. To account for the lack of WTC11 data available, we also collected all such datasets for H1-ESCs for inclusion in the predictive model. This resulted in 1,506 bigWig files for HepG2 cells; 1,206 files for K562 cells; and 277 files for WTC11/H1-ESCs (**Supplementary Table 5**). For each candidate element aside from controls, we computed the mean bigWig signal extracted from the corresponding region of the human genome using bigWigAverageOverBed^50^. All data was right-skewed, and was therefore log-transformed (*i.e.*, after adding a pseudocount of 0.1) to approximate a normal distribution. Finally, for each cell type, multiple replicates corresponding to the same “Experiment target” (**Supplementary Table 5**) were averaged to compute the consensus signal for each target in each cell type. This led to a total of 655 HepG2 features, 447 K562 features, and 122 WTC11/H1-ESC features considered by the models.

#### EnformerMPRA model features

For the large-scale libraries, Enformer^32^ was used to predict element activity in both orientations (*i.e.*, including adaptors in a fixed orientation to simulate the MPRA experiment), and the resulting 5,313 human predictions for each of the two orientations were averaged. For the joint library, Enformer was used to predict element activity in only the forward/sense orientation, and the resulting 5,313 human predictions were carried forward as features. As Enformer requires an input sequence length of 196,608 bp, all elements were extended with “N” padding on both cis and trans direction while keeping the element sequence centered.

#### Data pre-processing and model training

For each of the three large-scale libraries, the log_2_(RNA/DNA) scores for each element were averaged among both orientations in which the element was tested, and then randomly assigned to one of ten cross-validation folds. All predictive features (*i.e.*, biochemical features from the matched cell type, or all Enformer features) were z-score normalized to scale the features similarly. This enabled a direct comparison of coefficients among features derived from the resulting linear models. As described before^15, 19, 31^, for each regression task we optimized the *λ* regularization hyperparameter using 10-fold cross-validation, and then used the optimal value for *λ* to train ten lasso regression models, each on 9 of the 10 folds of data, to evaluate the performance of each model on the held-out fold. To evaluate the most relevant features selected, we trained a lasso regression model on the full dataset and visualized the 30 coefficients with the greatest magnitude. A similar strategy was used for data from the joint library tested in all three cell types, ensuring that the same element measured in different cell types was always assigned to the same fold.

#### Training MPRAnn

To derive sequence-based features we trained a simple convolutional neural network (CNN), implemented in tensorflow v2.6.2, with a total of 4 convolutional and 3 dense layers on the large-scale libraries. The complete model architecture is provided (**Extended Data Fig. 6a**). As input we used the 230 bp sequences including the adapters, one-hot encoded them and fit the mean log_2_(RNA/DNA) values from forward and reverse stands. We augmented the batches using the reverse complement of the 200 bp target sequence, while keeping the two 15 bp adapters fixed. To fit the model, we used a learning rate of 0.001, an early stopping criterion with patience of 10 on 100 epochs, and the Adam optimizer with a mean square error loss function. To make the results directly comparable to the lasso regression model, we trained MPRAnn using 10-fold cross-validation on the identical 10 folds of data as before. For each of the 10 folds, 9 models were trained using each of the remaining folds as a validation set. The final prediction was made on the held-out test set by averaging the predictions across the nine models. For training the joint library, we generated predictions for each of the cell types simultaneously using a multi-task framework. Here, no augmentation was performed because only forward sequences were tested in the MPRA. A snakemake pipeline implementing model training and prediction is available at https://github.com/visze/sequence_cnn_models.

#### Interpreting motifs identified by MPRAnn

As a step towards motif interpretation, *in silico* mutagenesis (ISM) scores were generated for all possible single nucleotide variants on each 200 bp sequence. Here, ISM scores were generated for sequences in each of the 10 CV test set folds, averaging predictions across the 9 corresponding models from the remaining training/validation sets. The average reference sequence prediction was then compared with that of the alternative sequence^31^. We then interrogated our ISM scores to identify the most pertinent motifs associated with changes in variant activity using TF-MoDISco-lite v2.0.4 (https://github.com/jmschrei/tfmodisco-lite), a more efficient version of TF-MoDISco^33^. The TF-MoDISco-lite algorithm was used with default settings and similar seqlet patterns were matched against JASPAR 2022 CORE vertebrate non-redundant database^53^ using Tomtom^36^.

#### In-silico mutagenesis scores on saturation mutagenesis MPRA elements

All variant effects of *F9*, *LDLR*, *PKLR*, and *SORT1* elements of the saturation mutagenesis MPRA^35^ were downloaded (https://kircherlab.bihealth.org/satMutMPRA) because they matched one of the three cell types (HepG2, K562, and WTC11) tested. Next, 1 bp deletions were removed from these datasets. ISM scores for all elements were then generated with MPRAnn using GRCh38 coordinates. Because most of the elements are longer than 200 bp element, the regions were tiled in 200 bp windows from the beginning with a step size of 150 bp, resulting in an overlap of 50 bp between neighboring windows. The predictions were averaged on overlapping windows. The missing 30 bp input of MPRAnn was achieved by adding the 15 bp adapter to both sides. Prediction and saturation mutagenesis data was compared on all variants with a minimum barcode support of 10 in the experiments.

#### Calculation of element specificity scores (ESSs)

To compute ESSs using activity scores from the joint library, log_2_(RNA/DNA) values from each cell line were first z-score transformed. Then an ESS for each element was computed by subtracting the element’s score in each cell type by the mean element score across cell types. A full table of ESSs is provided (**Supplementary Table 7**).

## Acknowledgments

We thank Jingjing Zhao for assistance with data deposition into the ENCODE portal as well as J. Stamatoyannopoulos’ and A. Kundaje’s group for sharing DNase data and peak calls, respectively. Research was supported in part on work supported under an NRSA NIH fellowship 5T32HL007093 (to V.A.); ASHBi, sponsored by the World Premier International Research Center Initiative (WPI), MEXT, Japan (to F.I.); DFG grant 464313370 (to M.S., M.D., and M.K.); and grants from the National Human Genome Research Institute (NHGRI) U24 HG009446 (to W.S.N.), and UM1HG009408 and UM1HG011966 (to J.S. and N.A.). J.S. is an investigator of the Howard Hughes Medical Institute.

## Author contributions

V.A. designed experiments, performed most computational analyses, and generated figures and tables. F.I. performed most MPRA experiments and wrote the associated methods sections. B.M., A.S., and Z.Z. helped with cloning and sequencing samples. M.S. trained and helped to interpret MPRAnn, and P.D. helped generate Enformer predictions and STREME results with supervision from M.K.. G.Y. and W.N. developed and performed the representative subset selection strategy. V.A. and N.A. wrote most of the paper with additional help from all authors. N.A. and J.S. supervised the study.

## Data and code availability

Raw sequencing data and processed files generated in this study are available for K562 (ENCODE IDs: ENCSR382BVV and ENCSR022GQD) and HepG2 (ENCODE IDs: ENCSR632EPR and ENCSR463IRX) cells. Additional raw data will be deposited during the review of this manuscript and is available from authors upon request. Code to train and interpret MPRAnn is available at https://github.com/visze/sequence_cnn_models.

## EXTENDED DATA FIGURES

**Extended Data Fig. 1:**
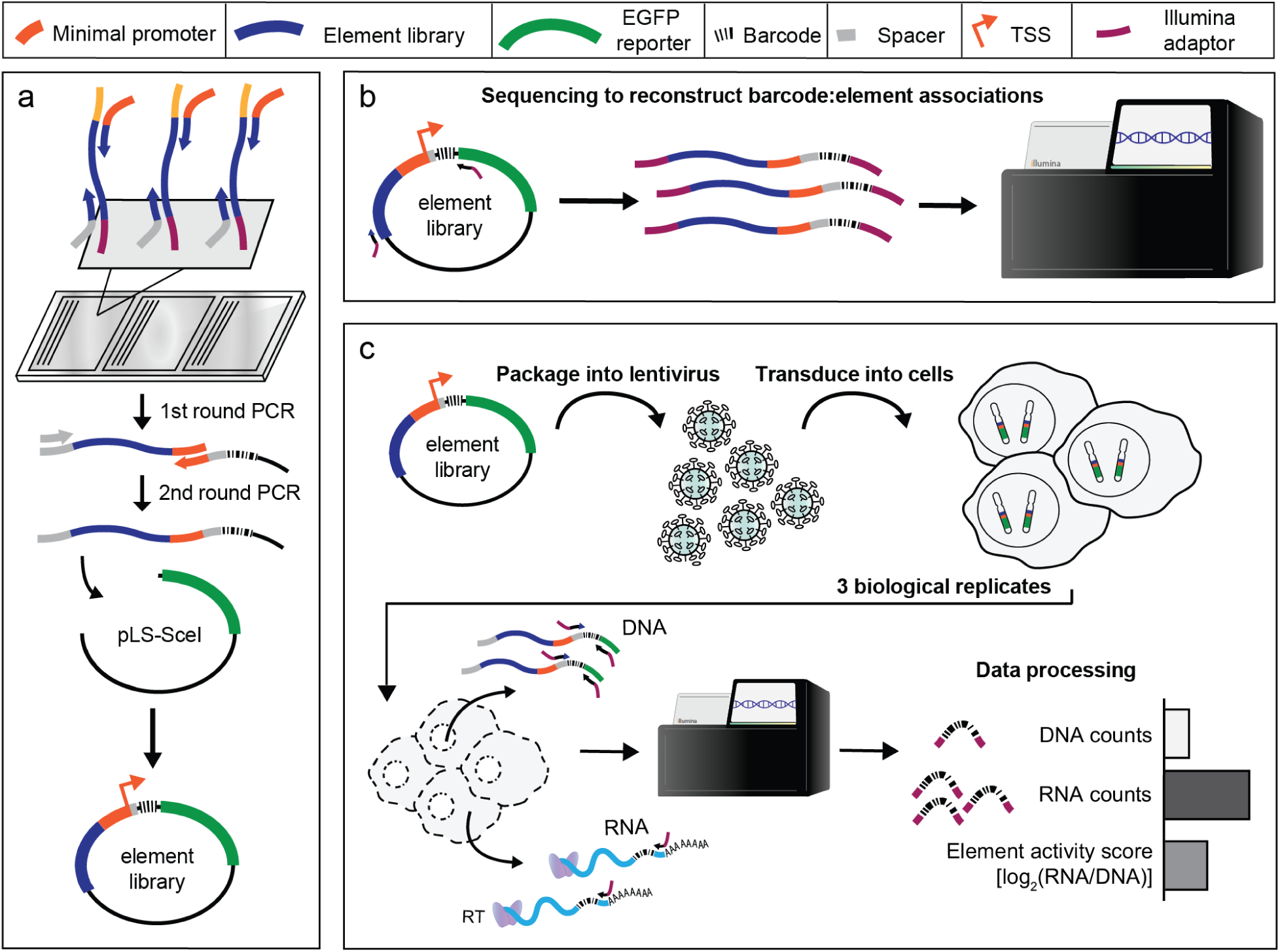
A next-generation lentiviral massively parallel reporter assay (lentiMPRA) strategy to measure the transcriptional regulatory activity of >6,000-240,000 enhancers simultaneously. **a,** Designed 230nt oligos corresponding to thousands of CREs are synthesized on an Agilent array. The 1st round of PCR adds on a minimal promoter, while the 2nd round of PCR adds random barcodes to these sequences. The library is then cloned into a pLS-SceI vector harboring an EGFP reporter to generate the final element library. **b,** The element-barcode fragments within the library are amplified by PCR and sequenced using an Illumina NextSeq instrument. This enables the reconstruction of which barcodes correspond to which enhancers. **c,** The enhancer library is packaged into lentiviruses and transduced into HepG2, K562, or WTC11 cells in a series of three replicates. Cells are grown in cultured medium for three days prior to the harvesting of RNA and DNA. Each RNA and DNA sample from each replicate is extracted, and barcodes are sequenced on an Illumina NextSeq instrument. Finally, DNA and RNA-derived barcodes are counted to compute a normalized activity score for each element in each replicate.

**Extended Data Fig. 2:**
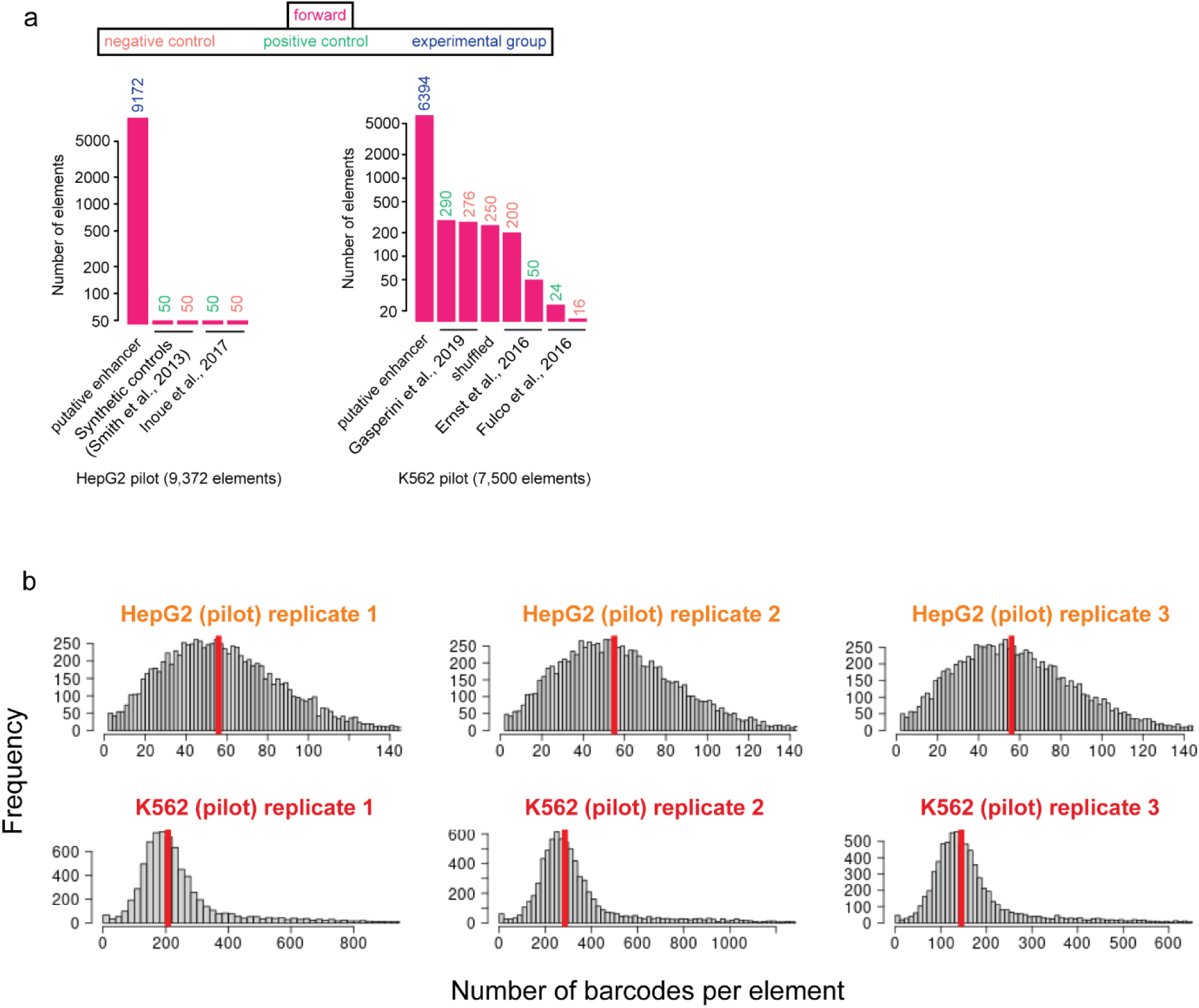
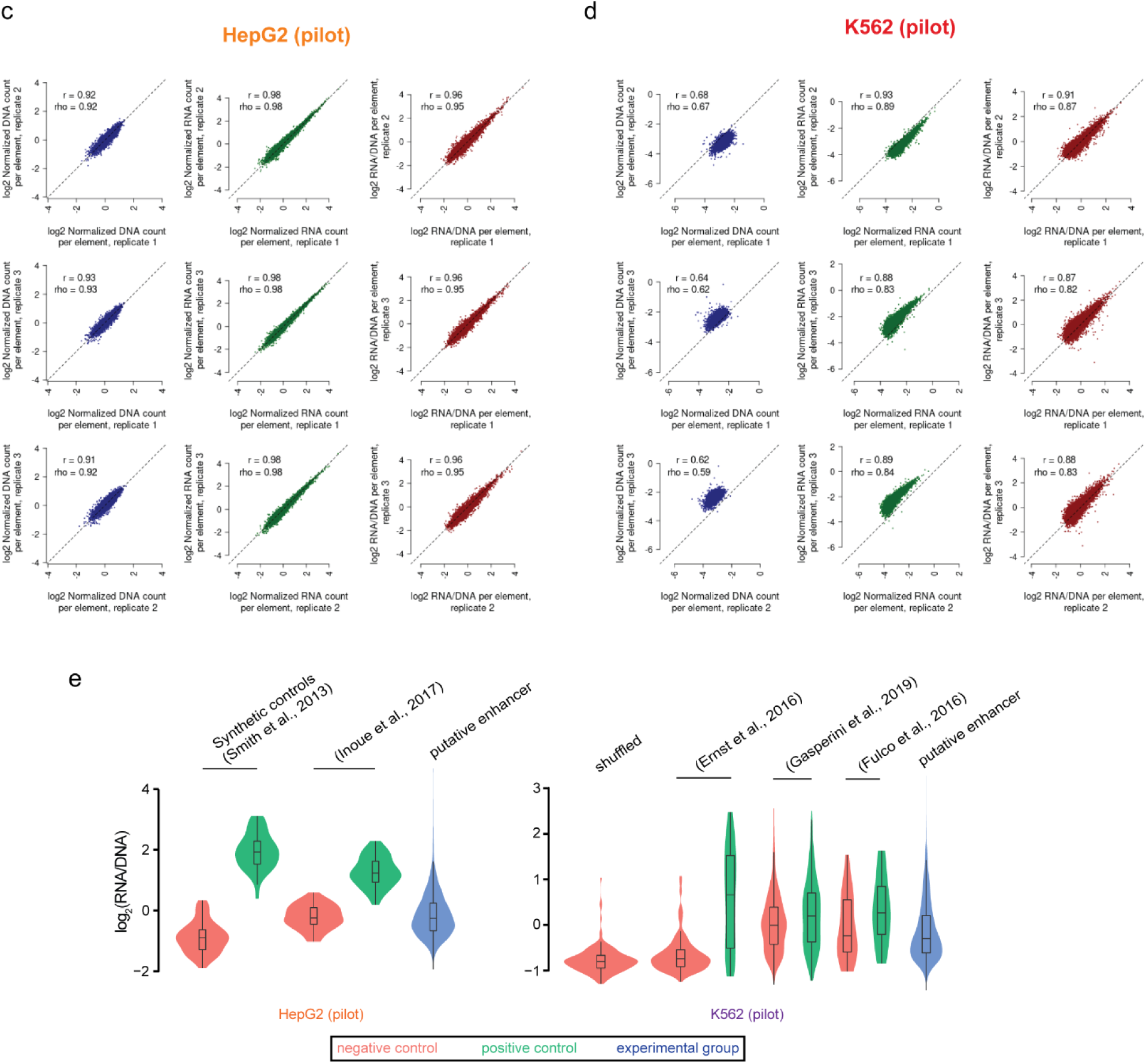
Design and quality control characteristics of two pilot MPRA libraries. **a,** Composition of the HepG2 and K562 pilot libraries. Thousands of putative enhancers, negative controls (dinucleotide shuffled sequences or elements lacking a signal from prior studies), and positive controls (elements with reported activity from prior studies) are included in each library^15, 21–24^. To maintain consistency with Fig. 1b, bars are colored according to orientation tested, with accompanying numbers indicating the number of elements tested in the category. Numbers are colored according to element type. **b,** Shown are histograms indicating the number of observed barcodes per element, for each of the three replicates and two pilot MPRA libraries. Shown with a vertical red line is the median number of barcodes per element. **c-d,** Shown are scatter plots displaying the relationship between observed DNA counts (blue), RNA counts (green), and RNA/DNA ratios (red) for all pairwise comparisons among replicates, for both the **(c)** HepG2 and **(d)** K562 pilot MPRA libraries. Also indicated is the Pearson (*r*) and Spearman (rho) correlation values. Candidate enhancers supported by fewer than 10 barcodes were filtered out prior to this analysis to reduce the impact of technical noise. **e,** Violin plots of element activity [measured as log_2_(RNA/DNA)] for putative enhancers, negative controls, and positive controls for each library.

**Extended Data Fig. 3:**
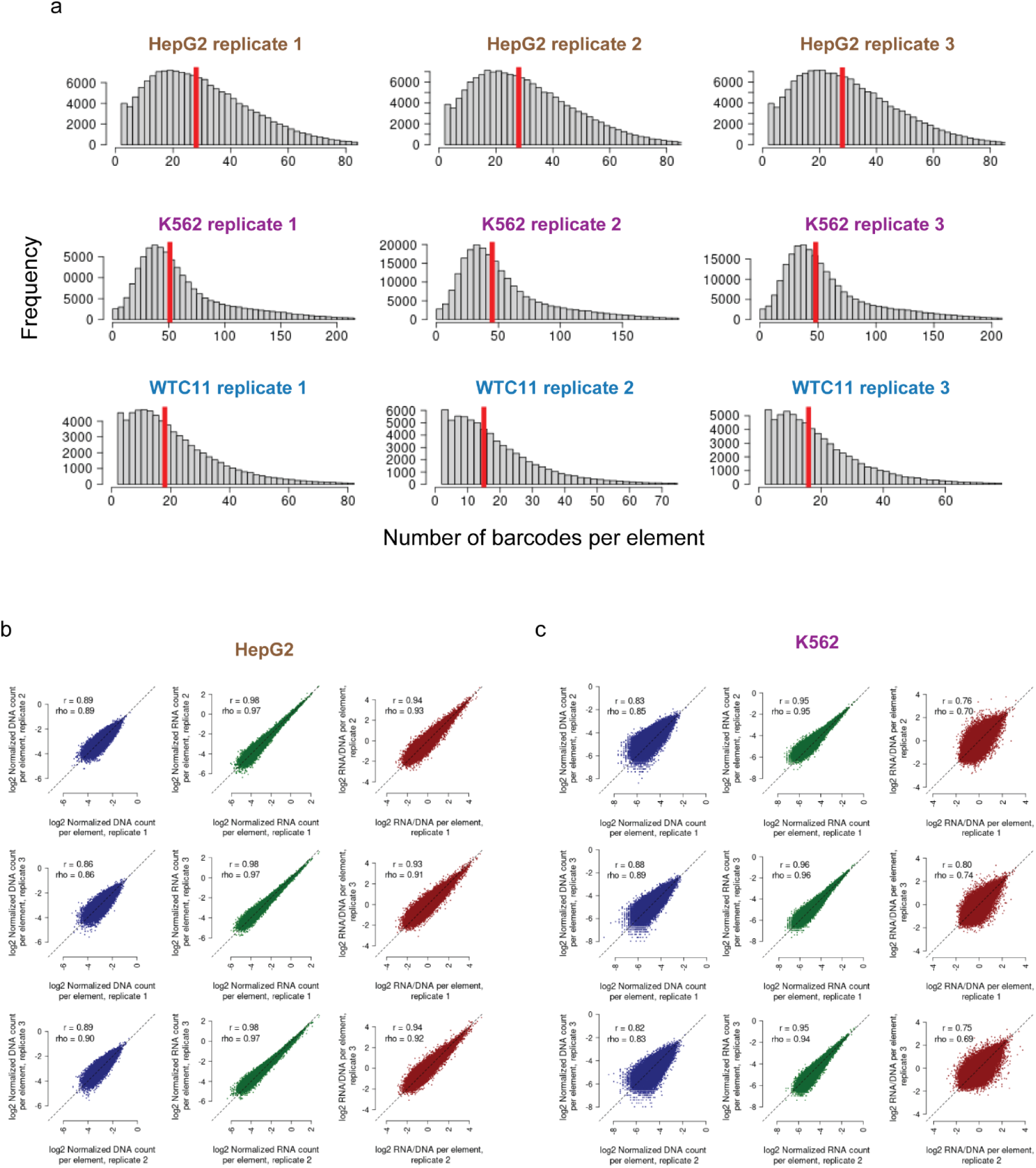
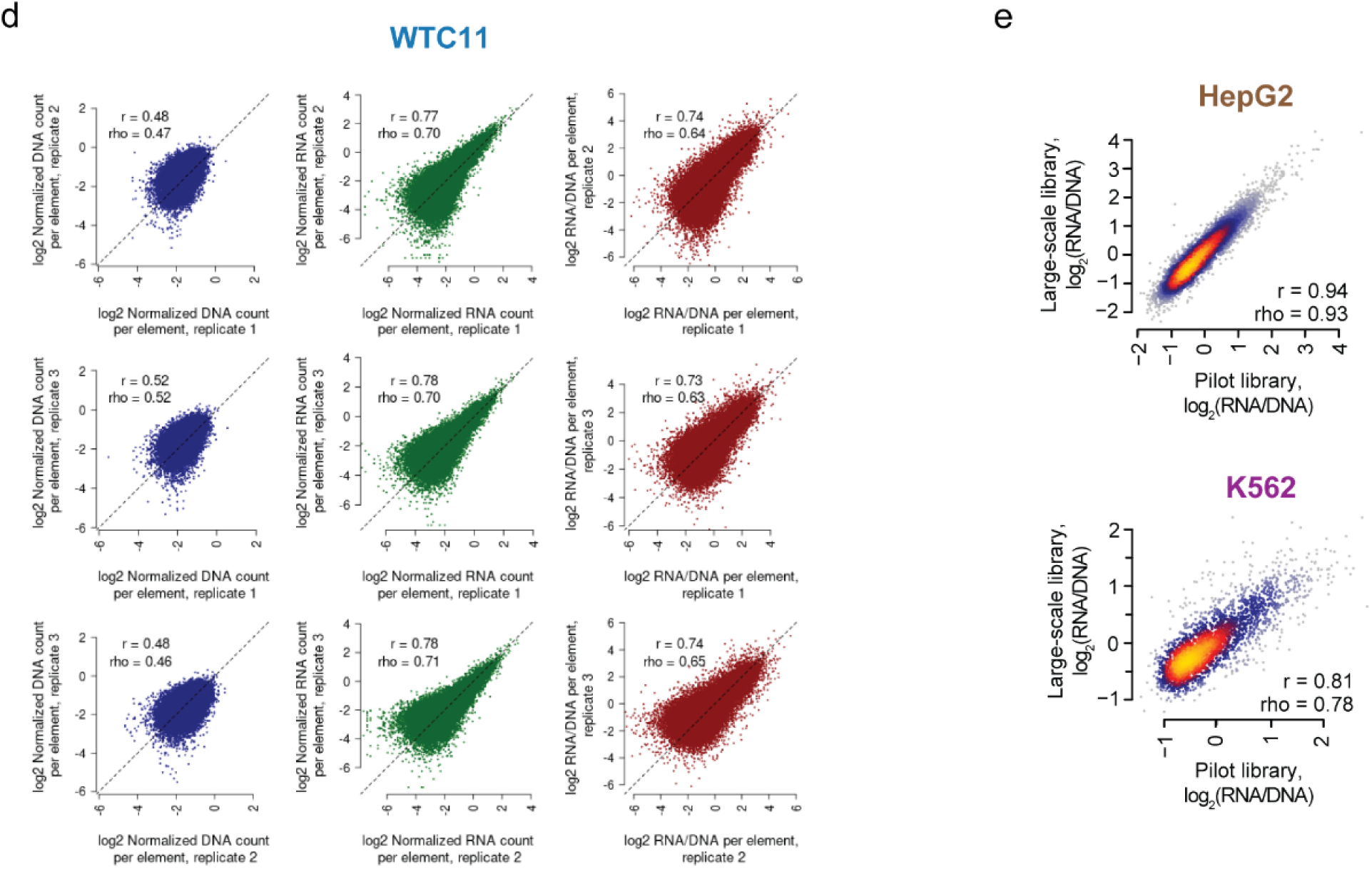
Quality control characteristics of the three large-scale MPRA libraries. **a,** Shown are histograms indicating the number of observed barcodes per element, for each of the three replicates and three large-scale MPRA libraries. Shown with a vertical red line is the median number of barcodes per element. **b-d,** Shown are scatter plots displaying the relationship between observed DNA counts (blue), RNA counts (green), and RNA/DNA ratios (red) for all pairwise comparisons among replicates, for the **(b)** HepG2, **(c)** K562, and **(d)** WTC11 large-scale MPRA libraries. Candidate elements supported by fewer than 10 barcodes were filtered out prior to this analysis to reduce the impact of technical noise. **e,** Scatter plots displaying the relationships between activity scores for the subset of elements common to both the pilot and large-scale libraries tested in HepG2 and K562 cells. Also indicated is the Pearson (*r*) and Spearman (rho) correlation values.

**Extended Data Fig. 4:**
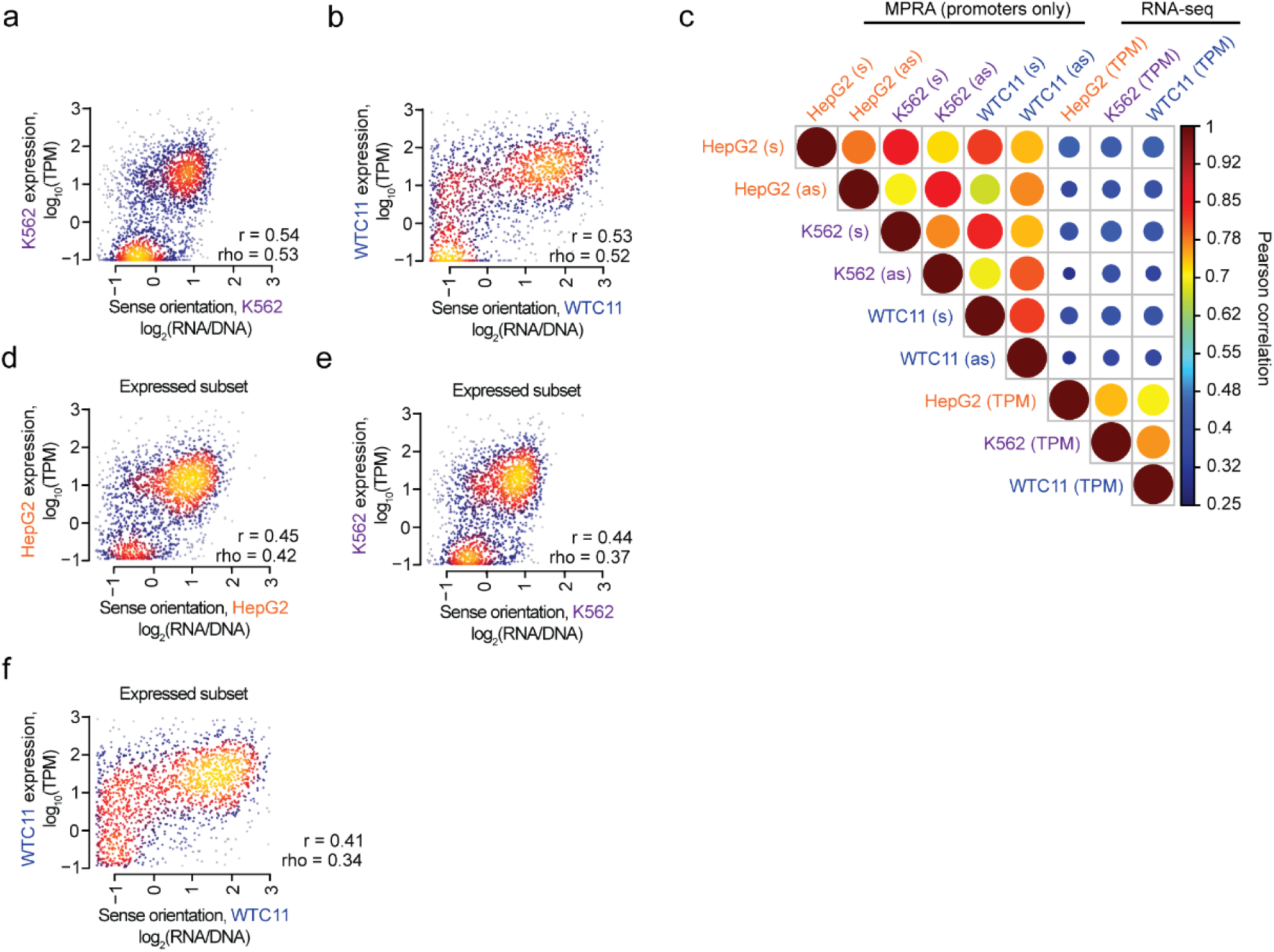
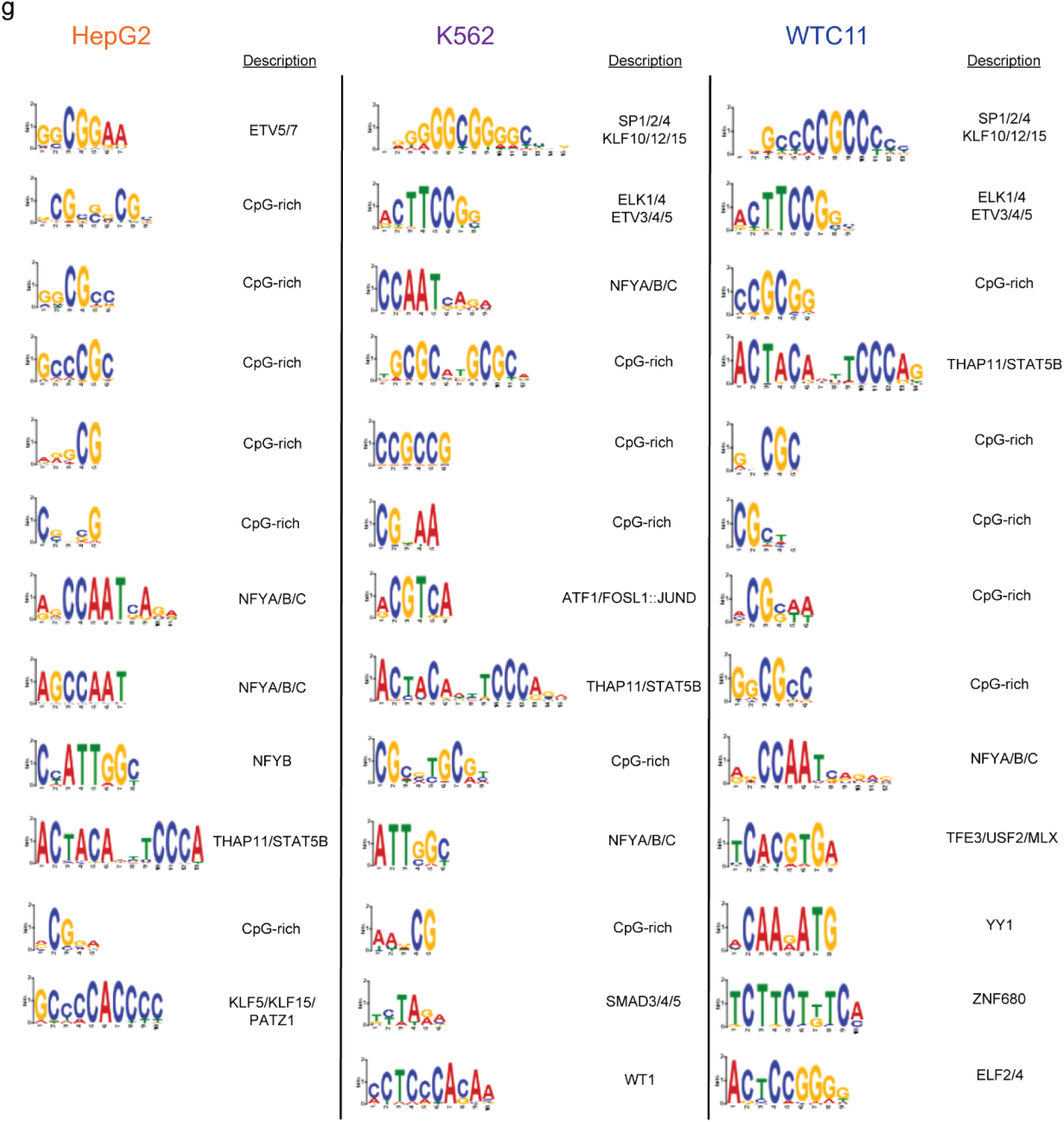
Properties of promoter activity in three cell types. **a-b,** Scatter plots of activity scores for sense-oriented promoters tested in the MPRA and endogenous gene expression levels for **(a)** K562 and **(b)** WTC11 cells. Expression levels follow a bimodal distribution. Also indicated are the Pearson (r) and Spearman (rho) correlation values. **c,** Upper triangular heatmap indicating the correlation between the sense (s) and antisense (as) orientations of promoters tested in the MPRA as well as endogenous gene expression levels measured in transcripts per million (TPM) using RNA-seq, filtered for the set of genes with detectable expression (*i.e.*, >0 TPM). The sizes of the circles are proportional to the Pearson correlations. **d-f,** Scatter plots of activity scores for sense-oriented promoters tested in the MPRA and endogenous gene expression levels for **(d)** HepG2, **(e)** K562, and **(f)** WTC11 cells, filtered for the set of genes with detectable expression (*i.e.*, >0 TPM). **g,** Set of motifs enriched in the top 1,000 most active vs. bottom 1,000 least active promoters (*i.e.*, as measured by large-scale MPRAs). Motifs were discovered by STREME^54^ for each of the three cell types evaluated, and annotated by Tomtom^36^ (*i.e.*, other than the set of CpG-rich motifs).

**Extended Data Fig. 5:**
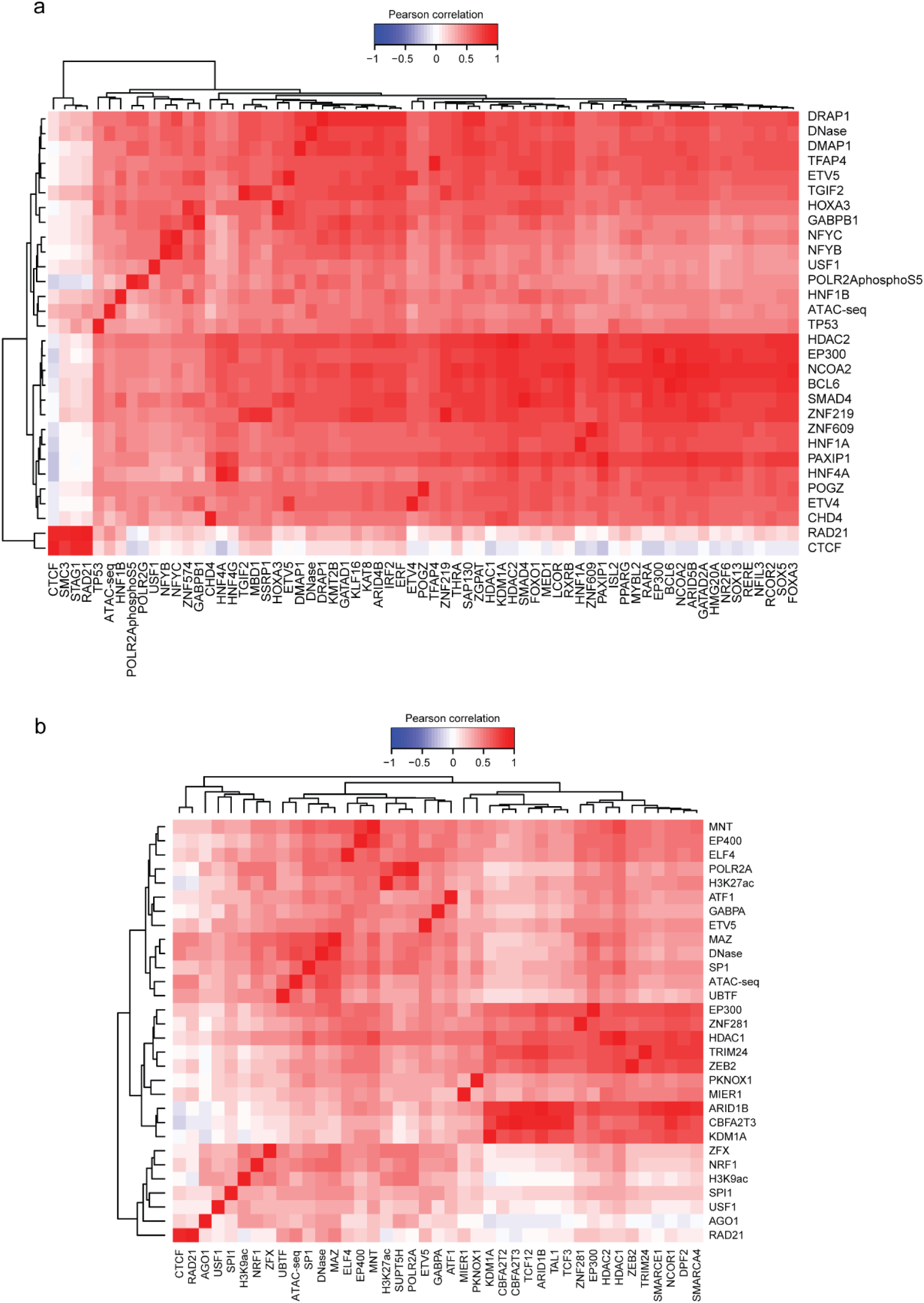
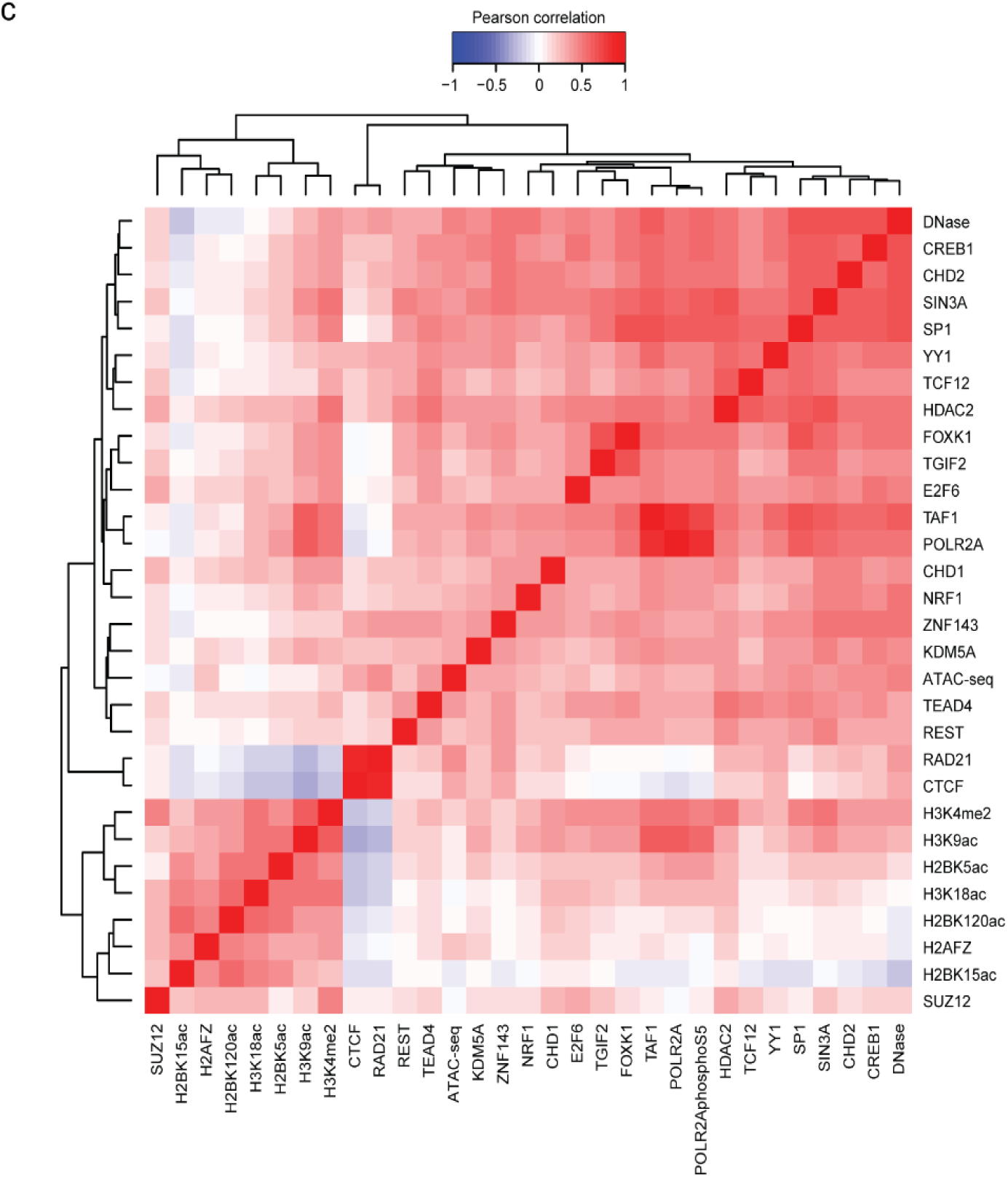
Alternative biochemical features that could explain element activity in large-scale MPRA libraries. **a-c,** Pearson correlation matrix between the top 30 features from **Fig. 3b**, shown as rows, and other features sharing a Pearson correlation either ≤ –0.8 or ≥ 0.8, shown as columns, for **(a)** HepG2, **(b)** K562, and **(c)** WTC11 cells. Hierarchical clustering was used to group features exhibiting similar correlation patterns.

**Extended Data Fig. 6:**
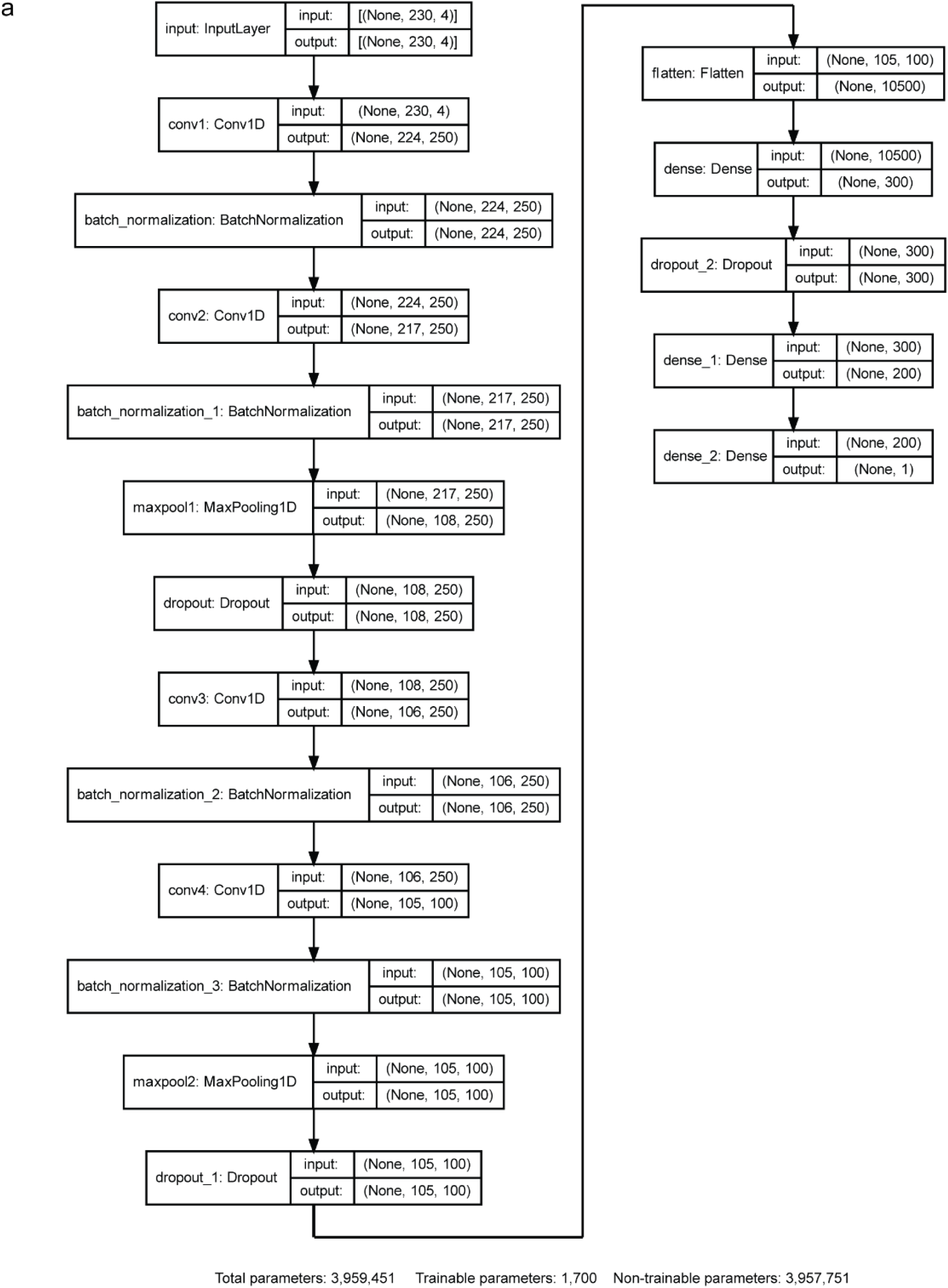
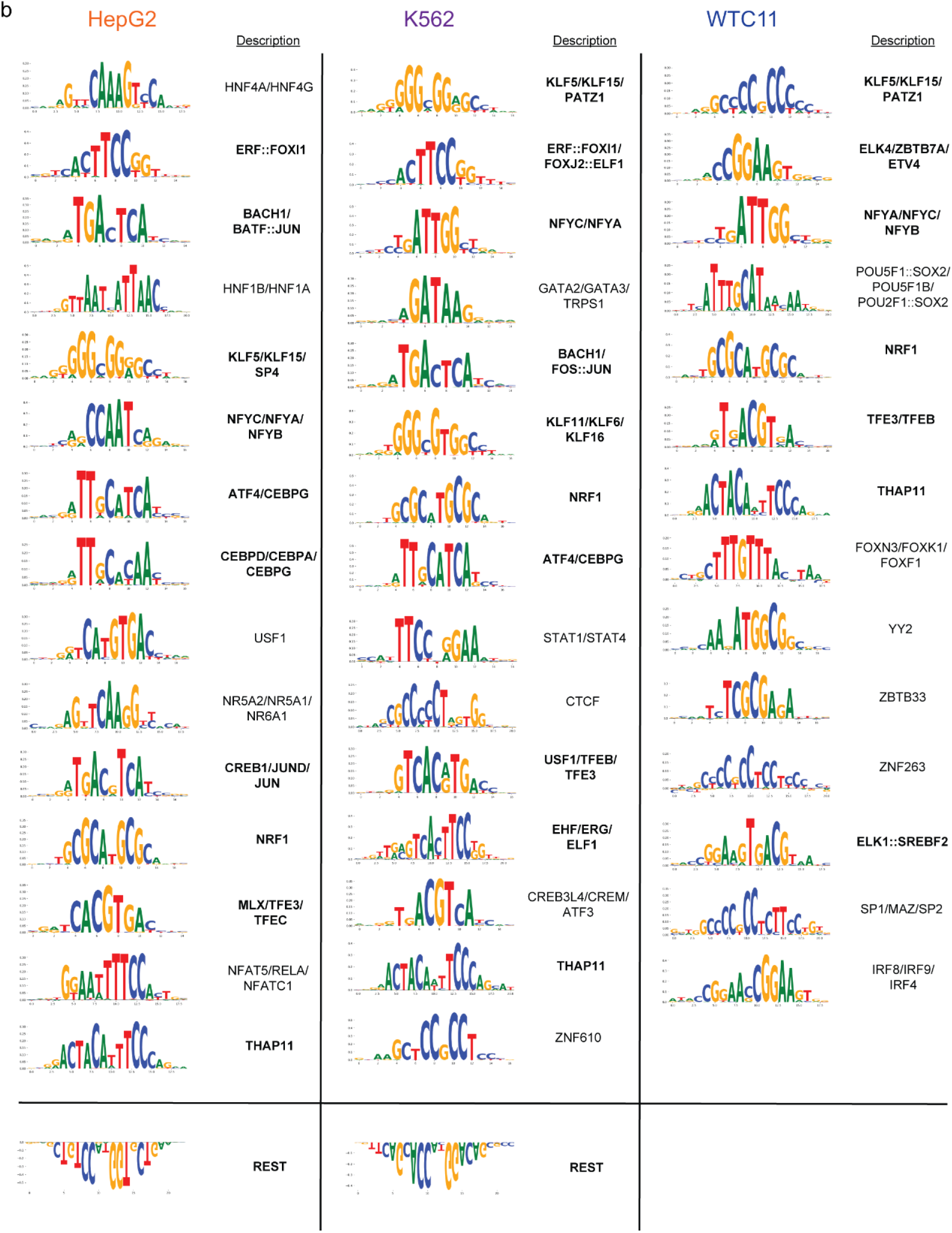
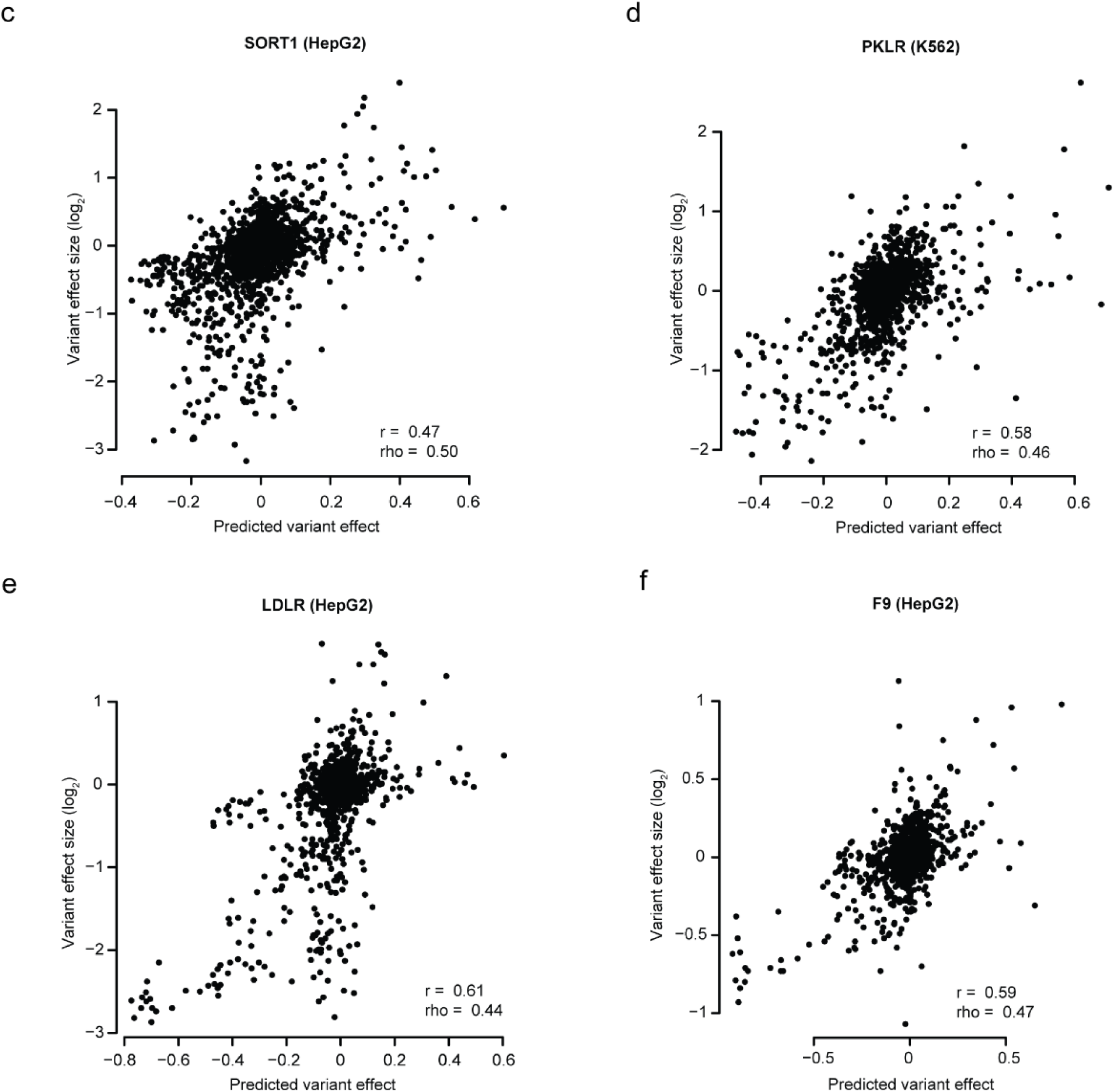
Architecture, interpretation, and performance of MPRAnn. **a,** Complete architecture of the MPRAnn model. Indicated for each layer is the layer name and dimensionality of the input and output matrices. **b,** Set of enriched motifs discovered by TF-MoDISco-lite^33^ for each of the three cell types evaluated. TFBSs associated with transcriptional inhibition (*e.g*., REST) are oriented upside down. TFBSs detected in at least two cell types (*i.e.*, likely bound to housekeeping TFs) are shown in bold. **c-f,** Scatter plots showing the correlation between predicted genetic variant effects by MPRAnn and observed variant effects, as detected in a saturation mutagenesis MPRA experiment testing the (**c**) *SORT1* enhancer, (**d)** *PKLR* promoter, (**e**) *LDLR* promoter, and (**f**) *F9* promoter^35^.

**Extended Data Fig. 7:**
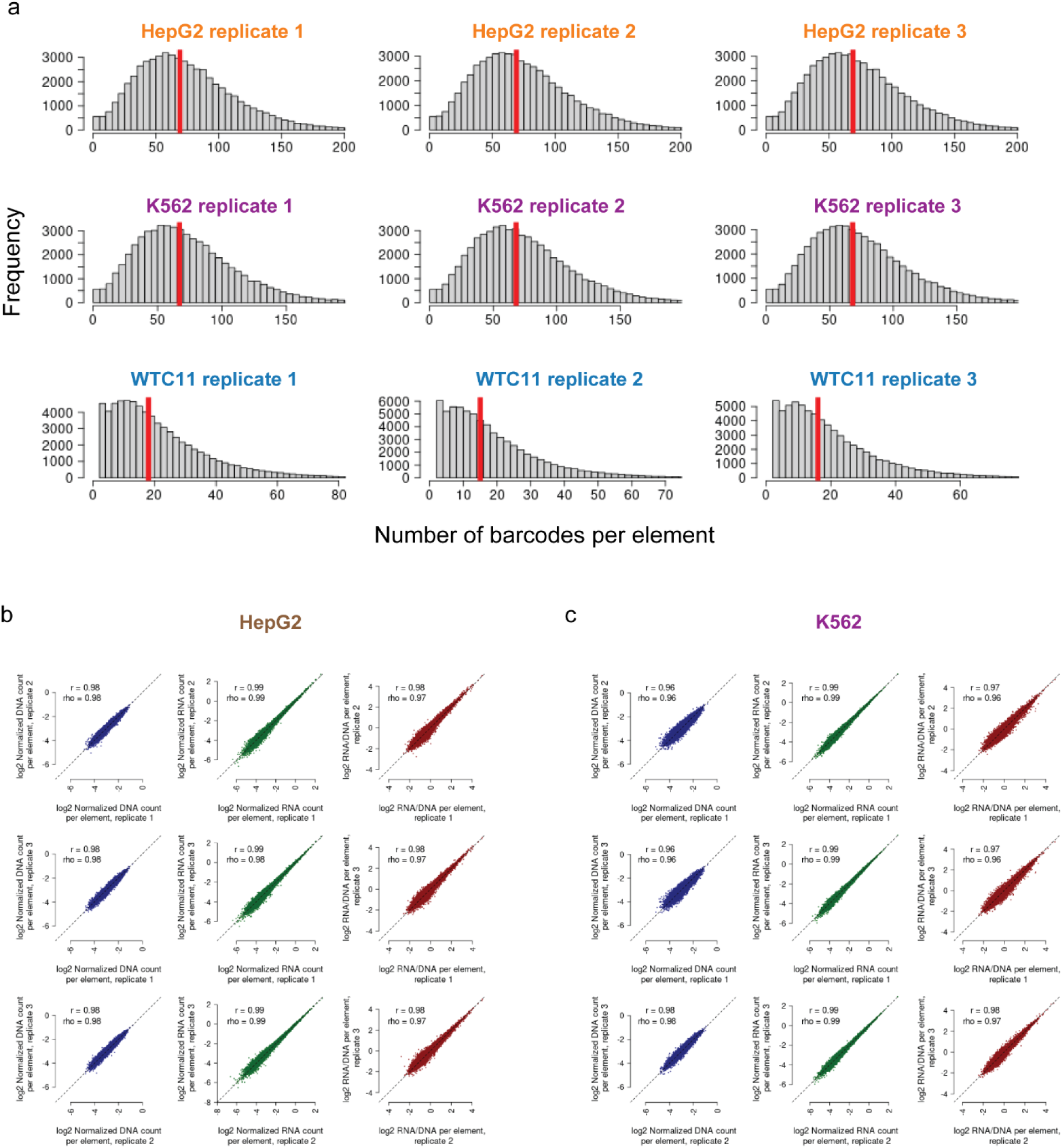
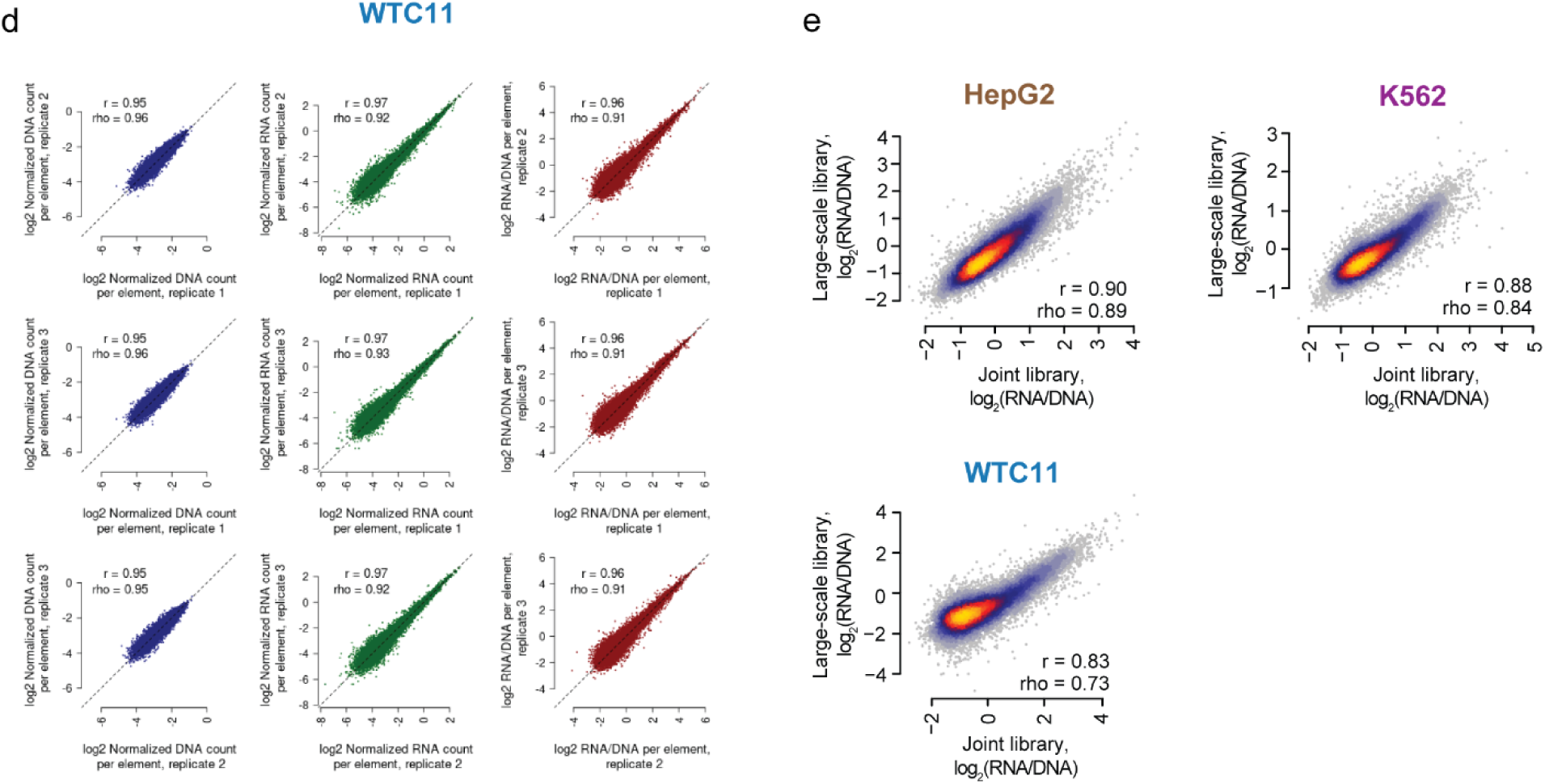
Quality control characteristics of the joint MPRA library. **a,** Histograms indicating the number of observed barcodes per element, for each of the three replicates and three cell types tested using the joint MPRA library. Shown with a vertical red line is the median number of barcodes per element. **b-d,** Scatter plots displaying the relationship between observed DNA counts (blue), RNA counts (green), and RNA/DNA ratios (red) for all pairwise comparisons among replicates, for the joint MPRA library tested in (**b**) HepG2, (**c**) K562, and (**d**) WTC11 cells. Candidate elements supported by fewer than 10 barcodes were filtered out prior to this analysis to reduce the impact of technical noise. **e,** Scatter plots displaying the relationships between activity scores for the subset of elements common to both the joint and large-scale MPRA libraries tested in HepG2, K562, and WTC11 cells. Also indicated are the Pearson (*r*) and Spearman (rho) correlation values.

**Extended Data Fig. 8:**
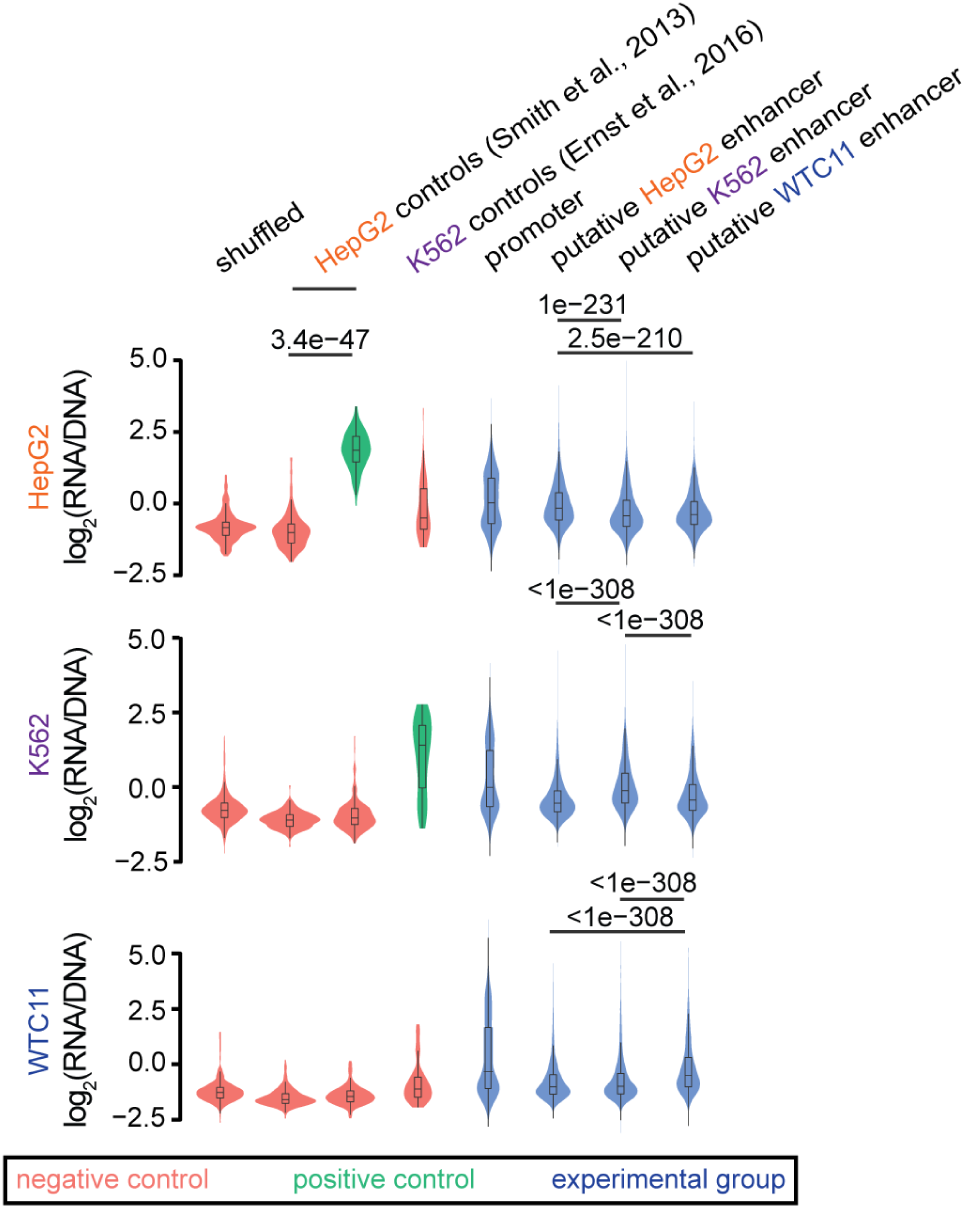
Violin plots of element activity [measured as log_2_(RNA/DNA)] in each of the three cell types for different element categories represented in the joint MPRA library shown in Fig. 5a. The difference between each pair of distributions tested was evaluated with a one-sided Wilcoxon rank-sum test, adjusted with a Bonferroni correction to account for the total number of hypothesis tests.

**Extended Data Fig. 9:**
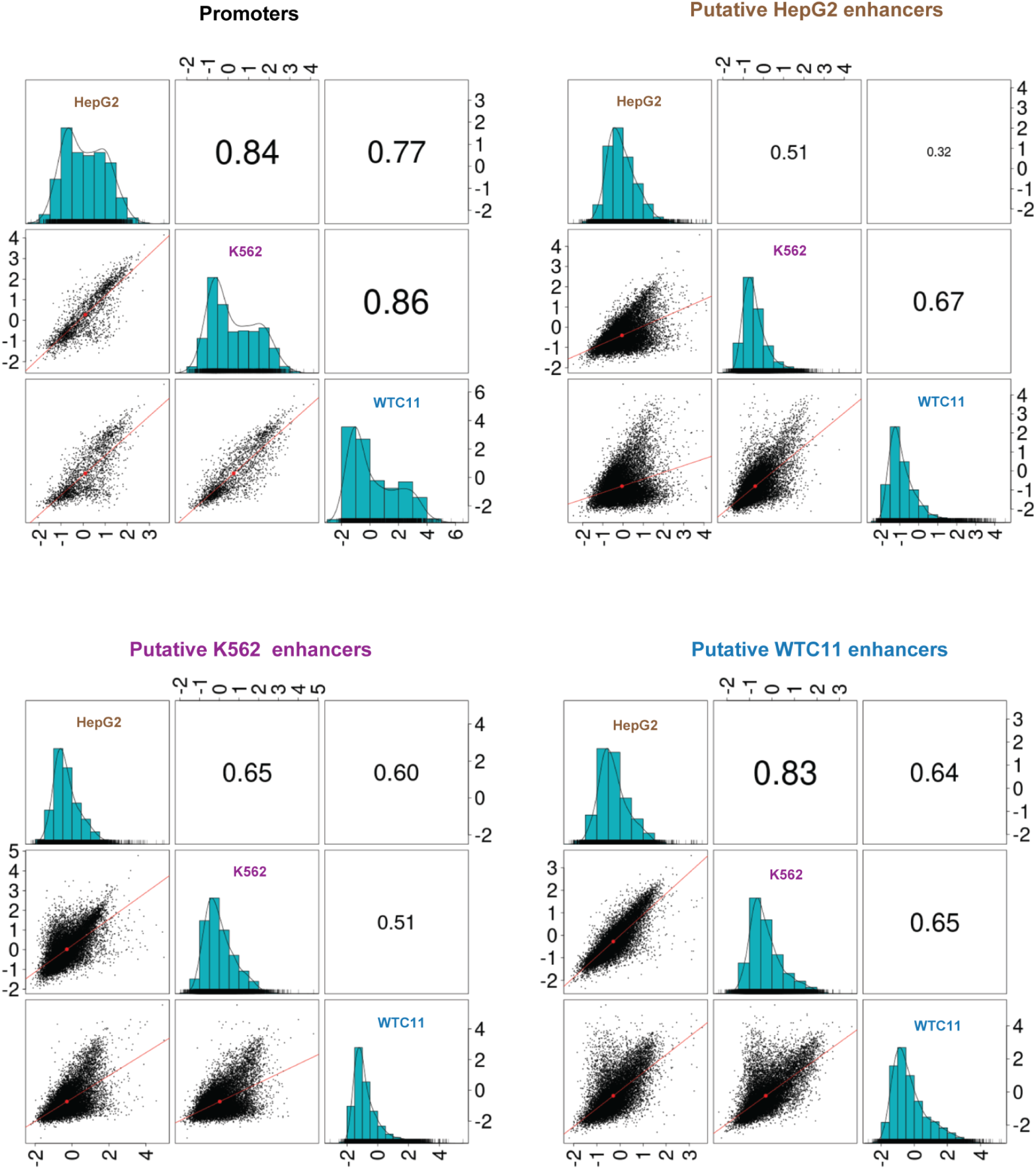
Comparison of promoter and enhancer activities from a joint MPRA library tested in three cell types. Scatter matrix displaying scatter plots corresponding to each of the three pairs of possible inter-cell-type comparisons (lower diagonal elements), for each of four element categories: i) protein-coding gene promoters, ii) putative enhancers selected from HepG2 cells, iii) putative enhancers selected from K562 cells, and iv) putative enhancers selected from WTC11 cells. Shown on the diagonal is a histogram of element activity scores [measured as log_2_(RNA/DNA)]. Also shown are Pearson correlation values among each pair of comparisons, with the size of the text proportional to the magnitude of the correlation coefficient (upper diagonal elements).

**Extended Data Fig. 10:**
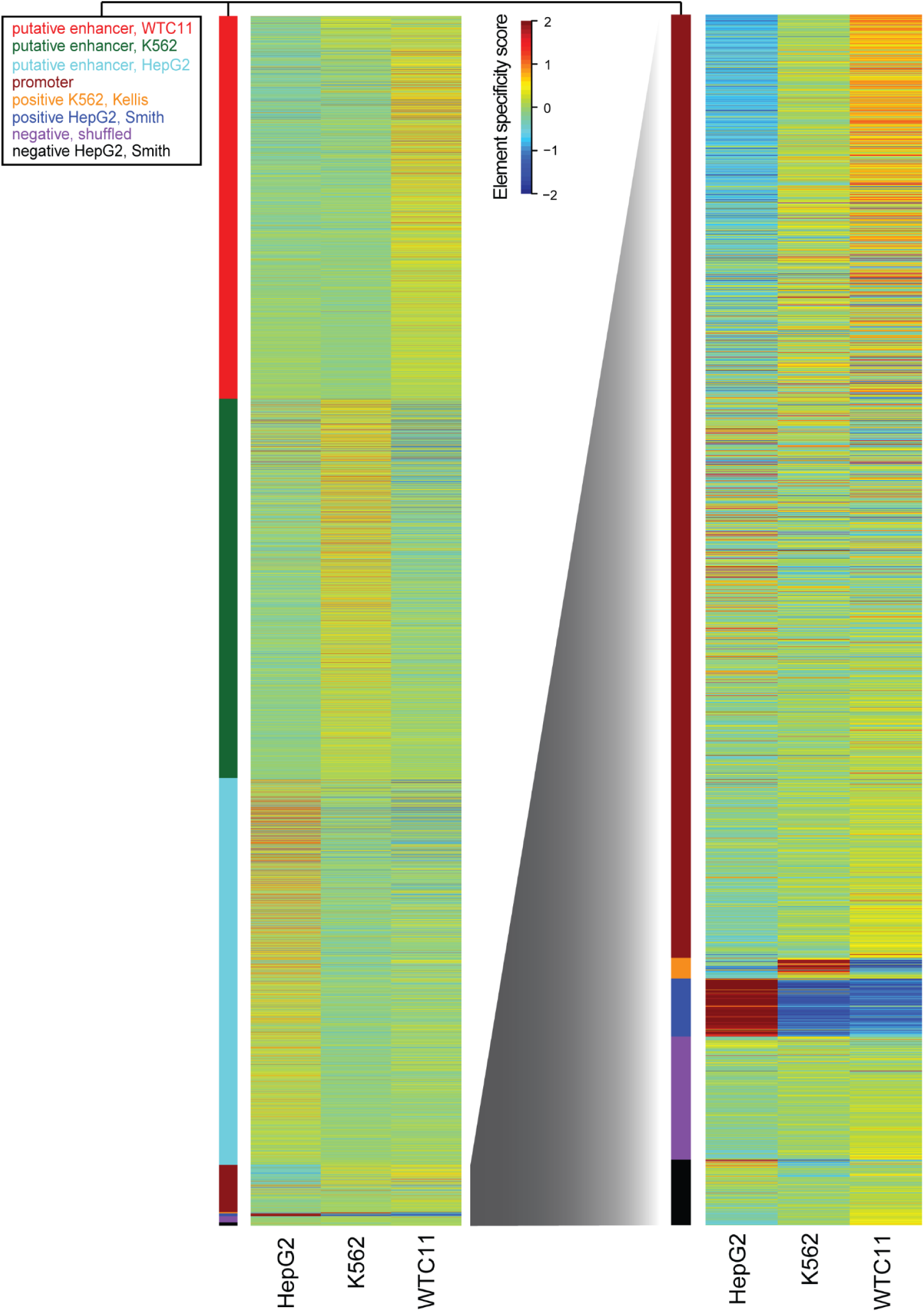
Comparison of element specificity scores derived from a joint MPRA library tested in three cell types. Heatmap of element specificity scores (*i.e.*, computed as the deviation of an element’s activity from its mean activity in all cell types). The heatmap shown on the right is a zoomed version of the heatmap on the left for elements other than putative enhancers. Elements are colored according to their category in the key provided.

**Extended Data Fig. 11:**
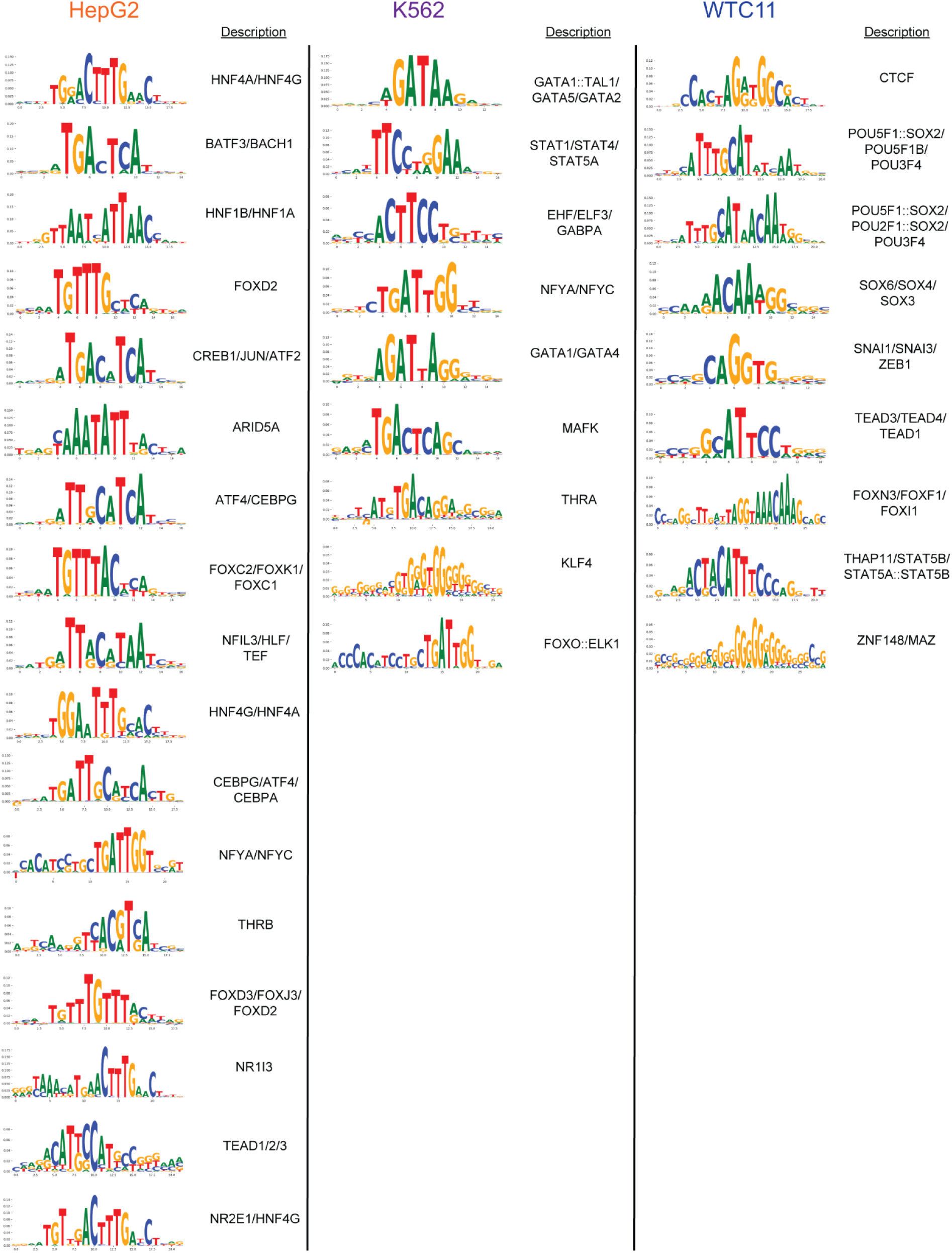
Interpretation of cell-type-specific motifs detected by MPRAnn. Set of enriched cell-type-specific motifs discovered by TF-MoDISco-lite^33^ for each of the three cell types evaluated on the joint MPRA library.

## SUPPLEMENTAL TABLES

**Supplementary Table 1.** Genomic coordinates, element categorization, and sequences for designed elements for the two pilot lentiMPRA assays from HepG2 and K562 cells.

**Supplementary Table 2.** Activity scores computed for each element for the two pilot lentiMPRA assays. Also provided are averaged activity scores across replicates as well as individual scores for each replicate alongside normalized DNA counts, normalized RNA counts, and the number of barcodes per element.

**Supplementary Table 3.** Genomic coordinates, element categorization, and sequences for designed elements for the three large-scale lentiMPRA assays.

**Supplementary Table 4.** Activity scores computed for each element for the three large-scale lentiMPRA assays. Also provided are averaged activity scores across replicates as well as individual scores for each replicate alongside normalized DNA counts, normalized RNA counts, and the number of barcodes per element. Activity scores for each orientation, for the subset of elements tested in both orientations, are also listed.

**Supplementary Table 5.** Summary of ENCODE samples compiled to compute biochemical features, with detailed metadata corresponding to the data source of origin, sample accession IDs, and additional factor-specific information. Also provided are pre-computed tables of the features used to train the biochemical lasso regression model.

**Supplementary Table 6.** Genomic coordinates, element categorization, and sequences for designed elements for the joint lentiMPRA library tested in three cell types.

**Supplementary Table 7.** Activity scores, and corresponding element specificity scores, for each element of the joint lentiMPRA library tested in each of the three cell types. Also provided are averaged activity scores across replicates as well as individual scores for each replicate alongside normalized DNA counts, normalized RNA counts, and the number of barcodes per element. Sheets summarizing the activity of each element across cell types as well as the derived element specificity scores are also included.

**Supplementary Table 8.** Primers used in this study.

